# Multi-omics-driven kinetic modeling reveals metabolic adaptations and vulnerabilities in BRCA1-deficient ovarian cancer

**DOI:** 10.1101/2025.10.22.682756

**Authors:** Ilias Toumpe, Maria Masid, Vassily Hatzimanikatis, Ljubisa Miskovic

## Abstract

*BRCA1*-deficient ovarian cancer cells undergo extensive metabolic reprogramming, yet the network-level dynamics underlying their proliferation and treatment response remain poorly resolved. Here, we construct large-scale multi-omics-driven kinetic model populations of ovarian cancer metabolism to track how tumor cells adapt to changes in nutrient use, energy production, and metabolite dynamics over time. Across *BRCA1* wild-type and mutant cells, these models expose distinct metabolic strategies shaped by transcriptional regulation and prioritize 28 enzyme-mediated vulnerabilities, including 24 linked to existing experimental or approved drugs and 4 previously uncharacterized targets in nucleotide and lipid synthesis. They further recapitulate a ceramide-linked metabolic stress signature shared across diverse chemotherapies. Mechanistic analysis traces the effects of *BRCA1* loss to transcription-factor-mediated shifts in enzyme activity, outlining regulatory routes for network-level rewiring. Beyond ovarian cancer, this framework offers a generalizable blueprint for predicting metabolic vulnerabilities, drug responses, and adaptive mechanisms across diverse cancer and metabolic disease contexts. By coupling dynamic metabolism to therapeutic prediction, it delivers actionable hypotheses for biomarker discovery, patient stratification, target prioritization, and precision metabolic medicine.

## Introduction

Altered metabolism is a hallmark of cancer.^1–3^ Cancer cells undergo extensive metabolic rewiring, which allows them to achieve elevated growth rates, survive anticancer treatments, and adapt to nutrient-deprived environments. Shared metabolic alterations occur across different tumor types, including the Warburg effect, enhanced biosynthesis, and increased energy demand.^4–6^ These conserved metabolic traits highlight metabolic vulnerabilities that can be therapeutically targeted to halt tumor proliferation.^7^ Indeed, some of the earliest chemotherapies, such as folate antagonists targeting nucleotide synthesis, exemplify how metabolic enzymes can be exploited for cancer treatment, inspiring the development of many subsequent anti-metabolite drugs.^8^

Epithelial ovarian cancer exemplifies how oncogenic mutations reprogram cellular metabolism. High-grade serous ovarian cancers frequently harbor germline or somatic *BRCA1* alterations, and *BRCA1*, beyond its role in DNA repair, modulates metabolism through transcriptional and regulatory networks.^9^ *BRCA1*-deficient (*BRCA1*^MUT^) cells show elevated glycolytic flux and altered mitochondrial function compared with wild-type (*BRCA1*^WT^). Most studies, however, assess these phenotypes via central-carbon proxies (glucose uptake, lactate secretion, oxygen consumption) that miss broader metabolic and regulatory changes.^10–12^ *BRCA1* also interacts with transcription factors governing nucleotide, lipid, and amino acid metabolism, indicating network-wide differences between *BRCA1*^MUT^ and *BRCA1*^WT^ tumors that extend well beyond glycolysis and oxidative phosphorylation. The lack of a metabolism-wide framework linking *BRCA1*-driven transcriptional changes to enzyme activities, fluxes, and metabolite levels limits mechanistic insight and therapeutic relevance.

Beyond this mechanistic gap, a broader translational challenge lies in turning metabolic insights into effective therapies. Despite promising preclinical data, clinical responses to metabolic interventions are highly context-dependent, often varying across tumor types, between subtypes of the same cancer, and even among closely related cell lines.^8,13,14^ These disparities reflect divergent metabolic states and microenvironmental constraints. In parallel, multiple chemotherapies induce a ceramide-centric sphingolipid stress signature across drug classes and tumor types, yet its origins, scope, and mechanistic basis across contexts remain unresolved.^15–18^ The basic challenge in fully understanding cancer cells’ responses to treatment is accurately monitoring drug-induced changes across the entire metabolic network, something that has proven challenging experimentally. Resolving this would uncover shared metabolic adaptations and biomarkers of response, reveal the rewiring that supports therapy resistance, and expose targets for rational, personalized therapy.

Computational efforts have sought to address these biological and translational gaps. Genome-scale metabolic models (GEMs) using flux balance analysis (FBA) have been applied to cancer metabolism.^19–36^ However, their steady-state nature prevents them from capturing time-resolved metabolite, enzyme, and regulatory dynamics that shape reprogramming and drug response.^31,32,37–43^ Consequently, they miss metabolic shifts underlying drug resistance^13^, toxicity^14,44^, and tumor-microenvironment interactions,^34,45,46^ where clinically relevant phenotypes emerge. These limitations highlight the need for frameworks that resolve transient metabolic and regulatory behavior.

Kinetic models meet this need by encoding enzyme mechanisms and regulation in ODEs^47^, enabling dynamic predictions and supporting diverse applications in human metabolism. ^23,38,47,48^ Smaller-scale studies have identified rate-limiting steps in glycolysis and central carbon pathways,^49–52^ while personalized red blood cell models illustrate how patient-specific enzyme activities can shape metabolic behavior.^38^ Capturing the full complexity of tumor metabolic reprogramming requires integrating multi-omics data into large- and near-genome-scale tissue- or patient-specific models that resolve transient metabolic behavior. A single kinetic model with a single parameter set offers only a narrow snapshot and cannot encompass the diversity observed across cell lines or among patients. In contrast, populations of kinetic models, varying in parameters and initial states, enable mapping of metabolic heterogeneity and reveal conserved regulatory patterns and shared, targetable vulnerabilities.

Here, we present curated large-scale kinetic models of ovarian cancer metabolism, built from multi-omics and physicochemical data for *BRCA1*^WT^ and *BRCA1*^MUT^ cell lines.^53^ These physiologically grounded model ensembles recover known metabolic vulnerabilities and identify four previously unrecognized cytostatic enzyme targets in nucleotide and sphingolipid metabolism. They resolve the network-wide transient responses to simulated drug treatments, capturing both established and previously uncharacterized metabolic shifts. The models reproduce a conserved ceramide-centered stress response across all simulated therapies and provide a mechanistic explanation for the metabolic fingerprint shared by diverse chemotherapeutic agents. By quantitatively reconciling the *BRCA1*^WT^ and *BRCA1*^MUT^ metabolic states, the models reveal a broad set of metabolic enzymes and fluxes under *BRCA1*-dependent regulation, linking multiple enzymes to upstream transcriptional programs shaped by *BRCA1* physiologies. Overall, these models provide a transferable mechanistic framework for probing metabolism across diverse disease settings, informing prognosis, treatment response, and the design of mechanism-informed drug combinations.

## Results

### Overview of the reconstruction framework for physiologically relevant kinetic models of cancer metabolism

We present a population of near-genome-scale kinetic models of cancer metabolism that capture the distinct physiologies of ovarian cancer cells with and without a *BRCA1* mutation and enable dynamic interrogation of enzyme-level vulnerabilities and network-wide adaptations. Kinetic model construction relies on a context-specific metabolic network derived from model reduction^54,55^ and, when available, integrated multi-omics data^37,40^. Here, we used and slightly extended a cancer-specific stoichiometric model^56^, derived from Recon3D^57^ through systematic reduction to ovarian cancer pathways and integrated with gene expression data from two isogenic cell lines (*BRCA1*^WT^ and *BRCA1*^MUT^) together with exofluxomic and exometabolomic datasets under thermodynamic constraints. The extension included incorporating measured adenylate levels (ATP, ADP, and AMP) and enforcing minimal enzymatic activity across all reactions to ensure global network connectivity. The resulting steady-state models capture distinct flux distributions consistent with differential gene activity and served as the scaffold for subsequent kinetic model construction (Figure 1A).

**Figure 1.**
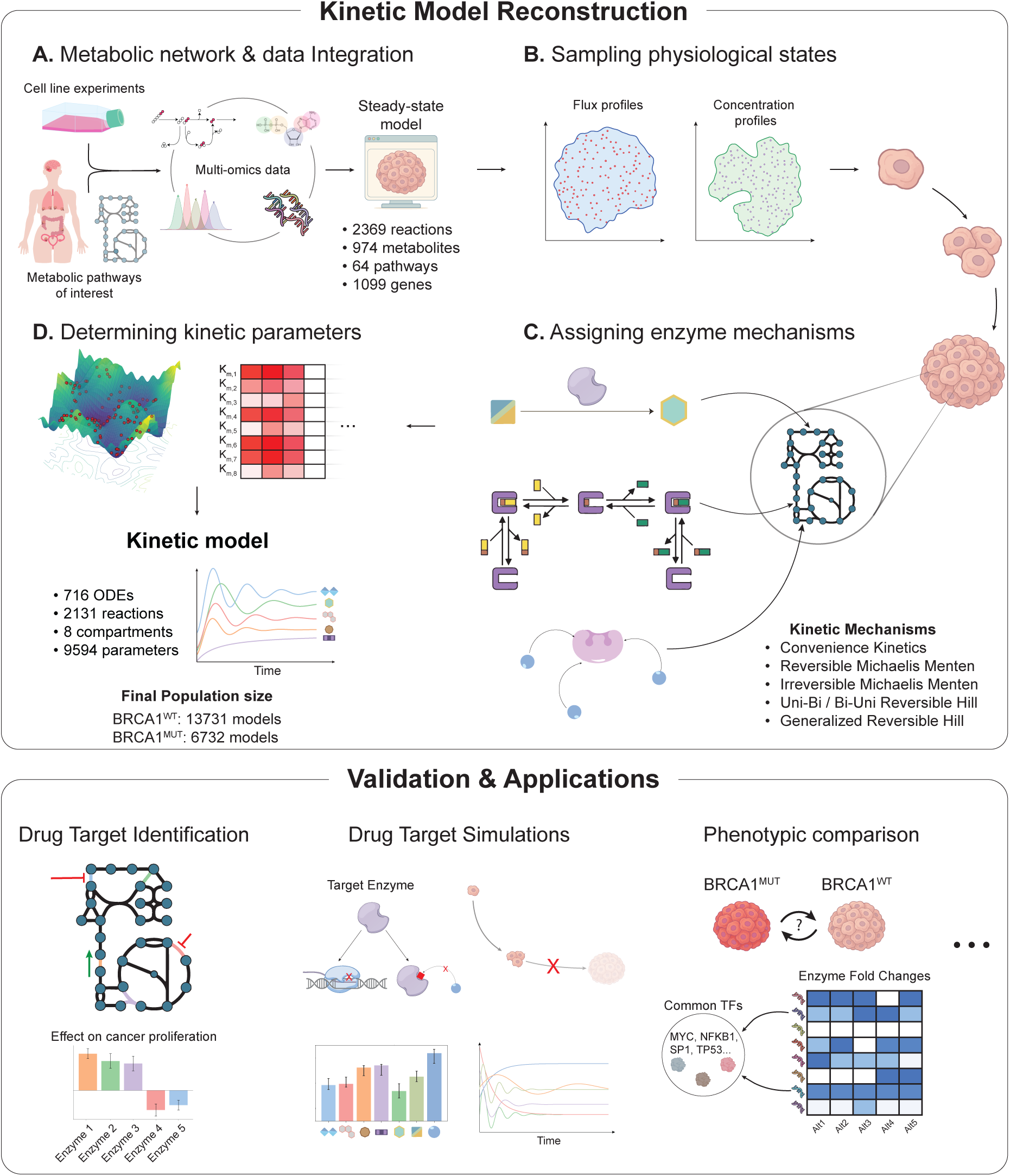
Overview of the reconstruction workflow, model characteristics, and applications of large-scale kinetic models of *BRCA1*^WT^ and *BRCA1*^MUT^ ovarian cancer metabolism. (A) The cancer-specific metabolic network was derived from the human genome-scale model Recon3D through a systematic reduction process. Transcriptomic, exometabolomic, and thermodynamic data were integrated to tailor the model to ovarian cancer physiology. (B) Sampling steady-state fluxes and metabolite concentrations produced physiologically relevant metabolic states that served as the basis for the kinetic models. (C) Each reaction was assigned a kinetic rate law mechanism, resulting in a model scaffold comprising hundreds of mass balances, rate laws, and kinetic parameters. (D) Parameter sets were sampled and filtered for physiological relevance, yielding populations of *BRCA1*^WT^ and *BRCA1*^MUT^ kinetic models that capture the observed metabolic diversity. These ensembles were used to identify drivers of proliferation, simulate drug administration, quantify pharmacodynamic responses and network-wide shifts, and link enzyme alterations to upstream transcription factors associated with *BRCA1*-dependent metabolic rewiring.

We next generated flux and concentration samples from each steady-state model to form a population of feasible metabolic states (Figure 1B). Sampling revealed multiple feasible metabolic phenotypes, highlighting the inherent metabolic diversity found among genetically similar cancer cells.^58,59^

Using the reduced metabolic network as the starting point, we constructed two large-scale kinetic model ensembles, one for *BRCA1*^WT^ and one for *BRCA1*^MUT^ ovarian cancer cells. Each model comprises more than 710 mass balances (ODEs) and more than 2,000 reactions governed by diverse kinetic rate laws that reflect the underlying enzymatic mechanisms (Figure 1C).^60^ In total, each model contains over 9,500 kinetic parameters. To parameterize the models, we used flux and concentration profiles obtained from the steady-state models and sampled kinetic parameters consistent with the observed metabolic behavior (Figure 1D, Methods). Model instances exhibiting non-physiological dynamics were discarded, ensuring that the final model populations capture biologically meaningful metabolic behavior beyond what steady-state models alone can reveal.

The curated model populations reproduce the metabolic diversity observed across cancer cells, providing physiologically grounded, dynamic representations of cancer cell metabolism. Validation against experimental data (Figure 1) confirms that the models provide a reliable representation of ovarian cancer metabolism. The curated *BRCA1*^WT^ and *BRCA1*^MUT^ model populations provide a platform to investigate (i) which enzymes are most critical for sustaining tumor growth and thus represent promising drug targets, (ii) how the metabolic network adapts over time when exposed to anticancer drugs, and (iii) which specific enzyme activity changes underlie metabolic differences between *BRCA1*^WT^ and *BRCA1*^MUT^ cells. Together, these applications establish the model ensembles as a systematic framework for exploring therapeutic vulnerabilities and regulatory rewiring in ovarian cancer metabolism.

### Drug target analysis predicts known and novel cytostatic targets

A central challenge in cancer research is identifying strategies to slow or halt tumor cell proliferation. Kinetic models of metabolism provide a quantitative means to assess how changes in enzymatic activity reshape the metabolic state and impact cancer cell growth. Using the *BRCA1*^WT^ and *BRCA1*^MUT^ kinetic model populations, we applied Metabolic Control Analysis (MCA)^61,62^ to quantify how changes in enzyme activity propagate through the network and alter reaction fluxes and metabolite concentrations. We focused on the control of cellular growth rate to identify enzymes whose activity most strongly determines proliferation, and thus represents potential metabolic drug targets. The MCA workflow yielded distributions of control coefficients, capturing the mean effect and variability of each enzyme’s impact on growth.

We identified the top 20 enzymes with the strongest effect on cancer proliferation in each of the two cell types (*BRCA1*^WT^ and *BRCA1*^MUT^), resulting in a combined set of 28 unique enzymes (Figure 2A). Whether enzymes exhibited positive or negative control over proliferation was generally consistent between *BRCA1*^WT^ and *BRCA1*^MUT^ models, although differences in magnitude revealed context-specific sensitivities. For example, most enzymes involved in central carbon metabolism have a greater effect on the growth rate of *BRCA1*^MUT^, indicating a stronger dependence on glycolytic activity (Figure 2A). On the other hand, mitochondrial complex I and multiple enzymes from lipid metabolism exert greater control over *BRCA1*^WT^ proliferation, reflecting a higher reliance on oxidative phosphorylation and lipid biosynthesis (Figure 2A). Most high-control enzymes have a positive effect on cancer cell proliferation, indicating that their inhibition would attenuate tumor growth, e.g., PGI, TMDS, and mitochondrial complex I. Conversely, a smaller subset exhibits negative control, suggesting that enhancing their activity could suppress cancer progression, including PPAP, NTD1, and MI1PP. Such inverse relationships have been observed in cancer metabolism. For instance, GLS2 is downregulated in hepatocellular carcinoma, and its overexpression has been shown to markedly impair tumor cell colony formation.^63^

**Figure 2.**
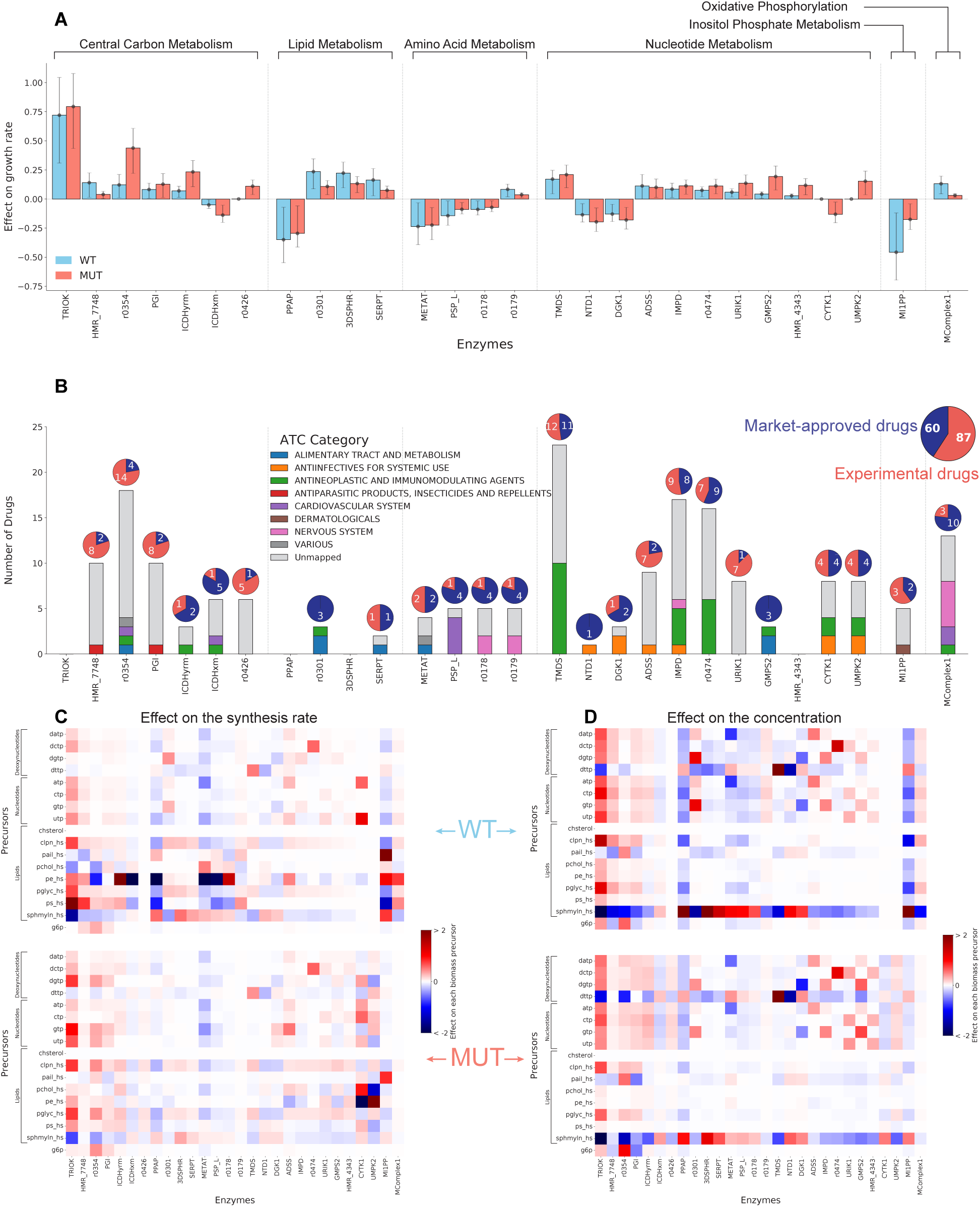
Metabolic Control Analysis of kinetic model populations identifies enzymes with strong control over cancer proliferation, their associated drug targets, and the impacts on biomass precursors. (A) Mean flux control coefficients (FCCs), quantifying the relative effect of a small change in enzyme activity on growth, for *BRCA1*^WT^ (blue) and *BRCA1*^MUT^ (red) kinetic model populations. Error bars denote the interquartile range (25th–75th percentiles). Displayed are the 28 enzymes with the highest absolute FCCs, grouped by metabolic subnetworks. Positive FCCs indicate that enzyme inhibition suppresses growth, and negative FCCs indicate that inhibition enhances proliferation. (B) Stacked bar chart showing the number of documented drugs targeting each of the 28 control-coefficient enzymes, classified by Anatomical Therapeutic Chemical (ATC) categories. Drug–target associations were curated from the DrugBank database. Pie charts illustrate the number of market-approved (blue) and experimental (red) drugs associated with each enzyme, with the larger pie chart displaying the totals across all targets. Heatmaps of (C) average flux control coefficients (FCC) and (D) concentration control coefficients (CCCs), quantifying how enzyme activity affects synthesis rates and metabolite concentrations, on biomass precursor classes: nucleotides, deoxynucleotides, lipids, and glucose-6-phosphate. Essential and nonessential amino acids are omitted due to minimal control. Color scale denotes the direction and magnitude of the CCCs (blue: negative effect; red: positive effect). Lists of enzyme and biomass precursor abbreviations are provided in Tables S1, S2.

To place these findings in the context of the broader cellular metabolic network, we mapped the functional roles of the 28 targets across major metabolic subsystems (Figure 2A), revealing their participation in anabolic (nucleotide and lipid biosynthesis), catabolic (glycolysis, TCA cycle, urea cycle), and bioenergetic (oxidative phosphorylation) pathways. These results align with therapeutic directions highlighted in recent reviews, which emphasize that targeting anabolic, catabolic, bioenergetic, and signaling processes represents a core strategy in cancer metabolism.^8,64^ Our models encompass the first three domains, while signaling remains beyond the scope of this study.

#### Validation with known drug targets and implications for repurposing

Having established a set of enzymes with strong control over cancer cell proliferation, we next examined which of these predicted targets are already associated with existing drugs and which represent novel predictions of the kinetic models. Of the 28 identified enzyme targets, 24 are linked to experimental or market-approved drugs in the U.S., Canada, and the EU, as cataloged in DrugBank^65^ (Figure 2B; Data S1). More importantly, 11 out of the 28 are targeted by at least one anticancer drug (Figure 2B). Notably, there are 20 known antimetabolites, which are structural analogs of natural metabolites that disrupt key metabolic pathways (ATC code L01B), and the models identified associated targets for 16 of these (Data S2). The remaining four include: (i) Carmofur, a fluorouracil derivative without documented targets and withdrawn from the market; (ii) one drug without associated enzymes in Recon3D; and (iii) two drugs whose targets appear in the reduced model but were not predicted here.

Many additional predicted targets correspond to drugs initially developed for non-cancer indications, with several already in clinical trials for the treatment of cancer. Overall, ∼60% of all documented drugs linked to the 28 targets are in experimental stages or clinical evaluation (Figure 2B), reflecting sustained therapeutic interest across multiple diseases, including cancer.

We also recorded each drug’s mode of action (inhibitory, activating, or antagonistic) (Data S1). Most act as inhibitors, consistent with our finding that most enzymes have a positive influence on growth. For example, thymidylate synthase (TMDS) is associated with 23 drugs, 10 of which act as inhibitors (Figure 2B, Data S1). In a few cases, however, drug effects diverged from model predictions. For example, IDH2 (ICDHxm) and CYTK1 were predicted to suppress growth when their activity increases (Figure 2A), yet both are inhibited by anticancer agents. However, it is worth noting that the documented anticancer drugs for IDH2 target mutant isoforms of the enzyme, which acquire neomorphic activity, producing the oncometabolite D-2-hydroxyglutarate.^66,67^ These inhibitors have been produced to selectively target mutant activity rather than wild-type IDH2. The apparent discrepancy thus reflects the specificity of current therapies for mutant IDH2 rather than a limitation of the modeling framework.

#### Novel targets and mechanistic insights

Among the enzymes highlighted by our models, most align with experimentally validated mechanisms, whereas the four with no associated drugs represent previously unexplored metabolic vulnerabilities. These novel candidates may offer promising avenues for future metabolic drug development, and we further evaluated their biological relevance and potential as therapeutic targets:

i. UPRT, a uracil phosphoribosyltransferase (HMR_4343). This enzyme catalyzes the salvage of uracil to UMP, a key precursor in pyrimidine nucleotide biosynthesis. In our model, inhibition of URPT reduces nucleotide pools and slows predicted growth rates, underscoring the enzyme’s potential role in sustaining rapid proliferation. Although the gene has been annotated in the human genome, its enzymatic activity has not been validated *in vitro*, and no studies have established a link between it and tumor biology.^68^ Nevertheless, pyrimidine metabolism is a well-established therapeutic axis, with multiple drugs targeting *de novo* or salvage pathways. The model positions URPT as a potential bottleneck in nucleotide supply and broadens the scope of established targeting strategies to an uncharacterized salvage enzyme, highlighting a gap where experimental validation could confirm a new metabolic vulnerability in cancer.
ii. TRIOK, a triokinase. TRIOK activity participates in the production of glyceraldehyde 3-phosphate. Perturbation of TRIOK alters flux through glycolytic and fructose metabolism pathways, affecting predicted growth rates, and suggests that cancer cells may rely on this enzyme when rewiring central carbon metabolism. While glycolysis and fructose metabolism are known to undergo rewiring in tumors, TRIOK itself has received little attention in cancer research.^69^ These findings raise the possibility that TRIOK supports metabolic flexibility in tumors, a role that remains to be tested experimentally.
iii. KDSR, a 3-dehydrosphinganine reductase (3DSPHR). Our model predicts that reduced KDSR activity slows growth by limiting the supply of structural sphingolipids required for membrane biosynthesis, leading to the accumulation of its substrate, 3-ketodihydrosphingosine. Blocking the activity of KDSR is toxic to cancer cells but not to normal cells^70^, providing a mechanistic link between the model prediction and selective cytotoxicity. Indeed, recent studies have demonstrated that many tumors rely on increased *de novo* sphingolipid biosynthesis to sustain their signaling and structural demands, and that targeting KDSR results in the buildup of 3-ketodihydrosphingosine and cancer cell death.^70,71^ Thus, while the model provides a direct metabolic explanation for slower growth, it also highlights the accumulation of 3-ketodihydrosphingosine as a measurable signature that can mechanistically connect to the downstream cytotoxic effects observed experimentally.
iv. PPAP, a phosphatidic acid phosphatase. Kinetic modeling predicts that increased PPAP activity suppresses growth by redirecting flux through lipid metabolism, specifically by converting phosphatidic acid to diacylglycerol and lowering the levels of the signaling lipid lysophosphatidic acid (LPA). This prediction aligns with experimental observations showing that phosphate phosphatase enzymes are significantly downregulated across multiple cancers, including ovarian cancer.^72–74^ Experimental studies have shown that introducing hLPP-3^75^ (which hydrolyzes phosphatidic acid, LPA, and sphingosine) or LPP-1^74^ into ovarian cancer cells reduces colony formation, promotes apoptosis, and inhibits tumor growth, suggesting that targeting LPA metabolism or signaling may be a promising therapeutic approach for ovarian cancer.^75^ Thus, while the model predicts growth inhibition through shifts in lipid metabolic balance, experimental evidence connects this mechanism to apoptosis and tumor suppression.

These findings demonstrate the model’s capacity to uncover both well-established and previously uncharacterized metabolic vulnerabilities. In some cases, the model directly predicts slower growth or altered metabolite levels, while in others, external experimental evidence links these metabolic changes to downstream phenotypes such as toxicity, apoptosis, or tumor suppression. Importantly, none of these targets were explicitly biased in the modeling process. Rather, their identification emerged naturally from the model architecture, kinetic formulation, and integrated multi-omics constraints.

#### Metabolic Enzyme Targets Disrupt Nucleotide and Lipid Supply

We interpreted the effect of each enzyme on cellular growth through its impact on the synthesis of biomass precursors: inhibiting an enzyme that depletes or limits the synthesis of a growth-limiting precursor is expected to decrease growth; conversely, increased precursor synthesis could promote it. In Recon3D, cellular growth is represented as the net synthesis of amino acids, nucleotides, deoxynucleotides, lipids, and glucose-6-phosphate. For each enzyme, we quantified its mean effect on the synthesis rate (Figure 2C) and intracellular concentration (Figure 2D) of each biomass precursor.

Most enzymes had minimal to no effect on either the concentration or the synthesis rate of amino acids, suggesting that cancer cells are robust to perturbations in amino acid metabolism. This finding aligns with prior reports^76,77^ showing that therapies targeting amino acid metabolism often have limited efficacy *in vivo*. Whereas in vitro assays typically probe isolated pathways, our kinetic models capture system-wide metabolic interactions that compensate for such perturbations, providing a mechanistic explanation for the reduced therapeutic response observed *in vivo*. Targeting the metabolism of essential amino acids is likely to cause systemic toxicity, while tumors can reroute fluxes around inhibitors of non-essential amino acid synthesis.^76^

In contrast, two precursors, sphingomyelin and deoxythymidine triphosphate (dTTP), were consistently sensitive to most enzyme targets for both physiologies (Figure 2C, 2D):

i. Sphingomyelin, which accounts for roughly 10–20% of cell membrane lipids, supports membrane rigidity and lipid raft formation through tight packing with cholesterol. It also serves as a source of signaling molecules such as ceramide and S1P, both regulators of apoptosis and proliferation.^78^ In our models, sphingomyelin is explicitly required in the biomass composition, ensuring that its contribution to growth is quantitatively represented. The models provide a non-obvious mechanistic insight by predicting that triokinase (TRIOK) and 3-dehydrosphinganine reductase (3DSPHR) exert the strongest control over sphingomyelin levels (Figure 2C, 2D). Specifically, increased TRIOK activity and decreased 3DSPHR activity are predicted to divert carbon away from sphingolipid metabolism, leading to sphingomyelin depletion.
ii. Deoxythymidine triphosphate (dTTP) is essential for DNA synthesis and repair, making it critical for rapidly proliferating cells. Cancer cells are particularly sensitive to disruptions in dTTP synthesis since limiting its availability causes uracil misincorporation, leading to DNA damage and cell death.^79^ Its *de novo* synthesis largely depends on a single enzyme, thymidylate synthase (TMDS), a well-established therapeutic target associated with many clinically approved anticancer drugs (as also predicted by the models; Figure 2B).^79^ The biomass formulation explicitly requires balanced pools of nucleotides, including dTTP, and the models predict that thymidylate synthase (TMDS) and 5’-nucleotidase (NTD1) are the most significant regulators of dTTP homeostasis (Figure 2C, 2D). TMDS is the only enzyme that produces *de novo* dTMP from dUMP, while NTD1 diverts dUMP away from dTTP production by converting it into deoxyuridine. Thus, the models show how dTTP balance can be controlled by both established and less obvious enzymes, reinforcing known strategies and revealing new vulnerabilities.

Overall, our framework successfully identified a set of enzymes with strong and specific control over cancer cell proliferation, many of which correspond to existing or emerging drug targets. Beyond growth control, the models uncovered systemic nodes that regulate the balance of key biomass precursors, providing mechanistic insight into how cellular homeostasis can be maintained or disrupted. In doing so, the models not only recapitulate well-validated therapeutic strategies but also highlight less obvious enzymes whose influence would not be apparent from single pathways alone. As this approach is extended to incorporate patient-specific omics data, it holds promise for guiding personalized interventions and precision oncology strategies.

### Simulations uncover transient metabolic shifts during cancer drug treatment

Despite decades of effort to target cancer metabolism, the mechanisms underlying many metabolic therapies remain poorly understood across the complex metabolic networks of both cancerous and healthy cells.^8^ Clinical efficacy is often inconsistent, and unintended side effects can emerge due to system-wide metabolic shifts.^80^ Dissecting these diverse outcomes is particularly challenging in human tumors, where dynamic microenvironments are experimentally complicated to probe. Kinetic models offer a systems-level solution to address these limitations by simulating drug administration and tracking how interventions impact metabolic fluxes and intracellular concentrations over time. Such simulations can reveal mechanisms and adaptive responses that are inaccessible through experiments alone.

We characterized intracellular metabolic responses to drug therapies by simulating pharmacological interventions against three functionally distinct metabolic enzyme targets in *BRCA1*^WT^ and *BRCA1*^MUT^ cancer models: (1) the hexokinases (HEX1, r0354, r0355), which catalyze the rate-limiting step of glycolysis and will serve as our primary case study; (2) thymidylate synthase (TMDS), a clinically validated anticancer target; and (3) triokinase (TRIOK), a novel growth-associated enzyme predicted by our models. All three were previously identified by Metabolic Control Analysis (MCA) as exerting strong control over cancer cell proliferation (Figure 2A).

#### Differential Drug Response Across Metabolic States

As a representative case, we first examined hexokinase inhibition in *BRCA1*^WT^ ovarian cancer (Figure 3), with analogous analyses performed across all targets and physiologies (Figure S1-S5). To capture distinct metabolic states and examine their differential responses to treatment, we selected a representative subset of kinetic models through stratified sampling based on their glycolytic and mitochondrial ATP production levels (Figures S6, S7). This sampling design mirrors patient stratification in clinical trials. It allows us to assess how cancer cells that regulate glycolytic and mitochondrial ATP production independently respond to targeted inhibitors.^5^ By doing so, the analysis captures metabolic heterogeneity, which is known to dictate therapeutic outcomes.^13,14^

**Figure 3.**
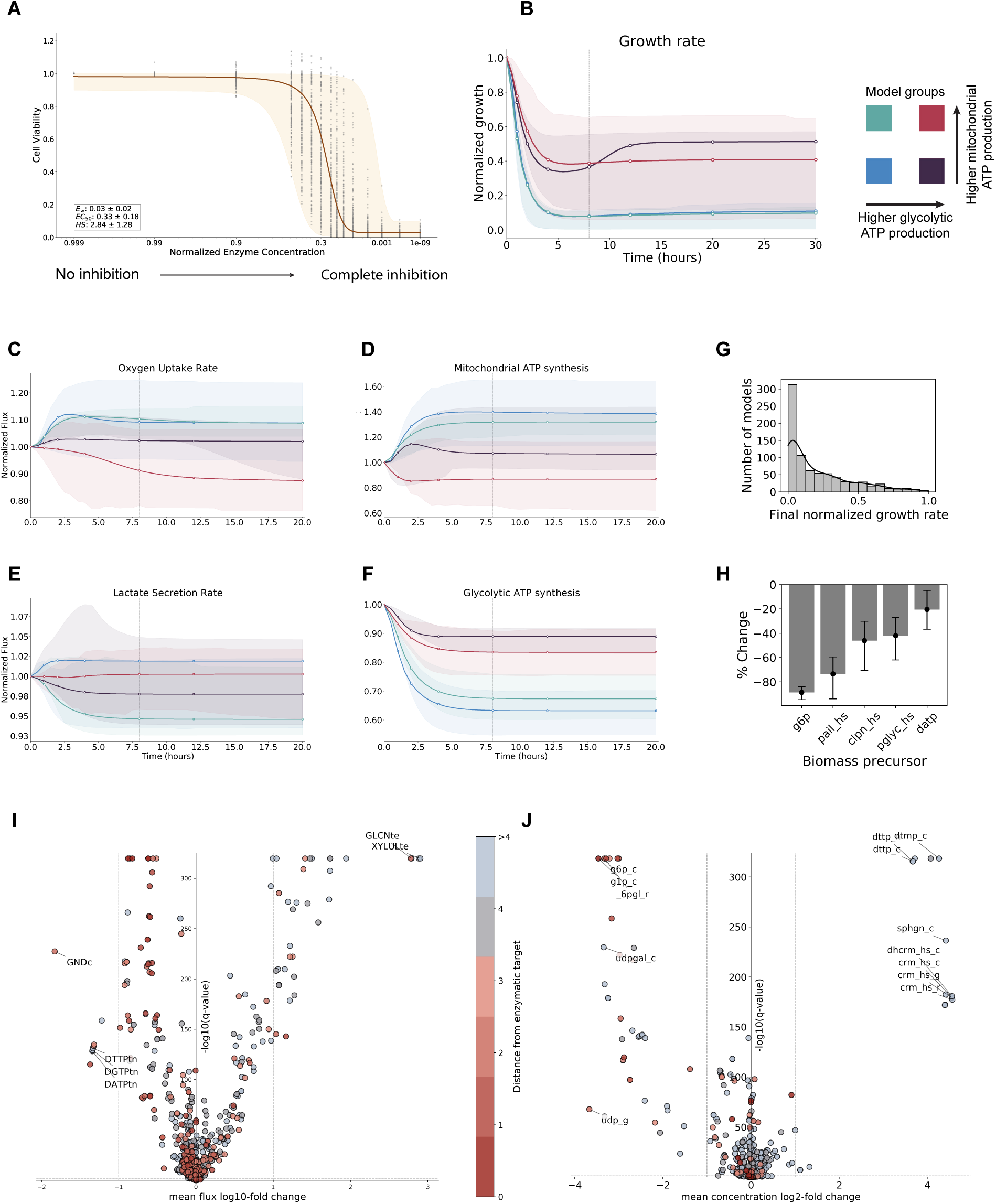
Hexokinase inhibition elicits dose-dependent growth arrest and network-wide metabolic adaptation in *BRCA1*^WT^ ovarian cancer models. (A) Dose–response curves of cell viability from 972 kinetic models, plotted against residual hexokinase activity. Despite variability in baseline states and parameters, all converge toward a common threshold: approximately two-thirds inhibition induces a 50% growth reduction (EC₅₀ = 0.33 ± 0.18). Grey dots represent individual model responses; orange-shaded ranges denote fitted logistic curves, with the average curve highlighted. (B) Transient growth trajectories following a 10-fold reduction in hexokinase activity, stratified by glycolytic vs. mitochondrial ATP production initial states (colored boxes). Shaded areas indicate interquartile ranges, and solid lines trace the model closest to each group’s mean. Transient responses of (C) oxygen consumption rate (OCR), (D) mitochondrial ATP synthesis, (E) lactate secretion rate, and (F) glycolytic ATP synthesis following the same inhibition. (G) Final growth rate distribution across the population following 10-fold inhibition. Nearly half of the models cease proliferation, underscoring pronounced heterogeneity in drug response. (H) Changes in biomass precursor pools after inhibition. Glucose-6-phosphate emerges as the principal bottleneck, accompanied by depletion of lipid species and dATP. Bars denote average percent changes, and the error ranges indicate the interquartile range for the five most affected precursors. (I-J) Volcano plots of steady-state changes in (I) reaction fluxes and (J) metabolite concentrations. Dots are color-coded by network distance from the inhibited hexokinases. Central carbon intermediates generally decrease, whereas several nucleotide and sphingolipid species accumulate, revealing pathway-specific rerouting upon glycolytic blockade.

We simulated simultaneous inhibition of all hexokinase isoforms by progressively reducing their enzymatic activity over an 8-hour window (Figure S8) and tracking the cancer cells’ metabolism until it reached a new metabolic state. The 8-hour timeframe was chosen as a conservative approximation of *in vivo* drug delivery, considering the limitations of tumor penetration and cellular uptake. In our framework, drug action was modeled by directly reducing the enzyme activity (Figure S8). However, if the mechanism of action is known, more mechanistic pharmacokinetic and pharmacodynamic (PK/PD) descriptions could be integrated, enabling simulation of competitive inhibition, allosteric modulation, or other regulatory effects.

Dose–response curves were generated for all selected models to assess cell viability, measured as changes in final growth rate across progressively reduced enzyme activity (Figure 3A). Because the models differ in their kinetic parameters and in the initial metabolic states at which inhibition is applied, the simulated responses varied substantially, mirroring the diversity of patient-to-patient physiology. Nevertheless, a clear pattern emerged: modest reductions in hexokinase activity had a limited impact, while complete inhibition reliably suppressed proliferation across all models. Logistic regression analysis was used to derive pharmacodynamic metrics such as EC_50_ (as performed in other studies^81^), revealing that, on average, a two-thirds reduction in enzyme activity was required to achieve 50% growth inhibition (EC_50_ = 0.33 ± 0.18). These simulations provide a mechanistic estimate of the degree of inhibition required to effectively impair cancer proliferation.

To investigate how cells adapt at the network level, we examined the effects of a 10-fold reduction in hexokinase activity on both fluxes and metabolite concentrations. The resulting responses varied across models with different initial glycolytic and mitochondrial activities, as shown in the apparent differences in final growth rates (Figure 3B) and internal metabolic states (Figure 3C-F). Models with lower glycolytic flux were generally more sensitive, showing reduced proliferation regardless of mitochondrial ATP output. Some models compensated by increasing the oxygen consumption rate (OCR) (Figure 3C) and mitochondrial ATP production (Figure 3D), while lactate secretion rate remained largely unchanged for all models (Figure 3E), even if glycolytic ATP generation decreased (Figure 3F). This compensatory increase in mitochondrial respiration following suppression of glycolysis has also been observed in cancer cells subjected to hexokinase inhibition.^82,83^ These results confirm that distinct metabolic states exhibit differential responses to the same enzymatic inhibition.

Population-level analysis showed that nearly half of the models ceased proliferation entirely (Figure 3D). Because the simulations track all intracellular metabolite concentrations and fluxes, we identified which biomass precursors were most depleted (Figure 3E). Glucose-6-phosphate was the principal bottleneck, consistent with its position as the direct product of the hexokinase-catalyzed reactions. Lipid species and dATP were also depleted, indicating growth arrest from the combined shortage of multiple precursors. This contrasts with the TMDS inhibition, where only dTTP was significantly depleted (Figures S1 and S4).

We next examined how hexokinase inhibition propagates across the metabolic network by comparing the initial and final metabolic states through the fold changes of fluxes and concentrations (Figures 3I, J). Central carbon intermediates generally declined, while several nucleotides and sphingolipid-related metabolites accumulated, revealing broad metabolic reprogramming (Figure 3J). Flux changes followed a similar pattern, with substantial decreases immediately downstream of hexokinase (one to four reaction steps) and a selective reduction in distal reaction fluxes, reflecting adaptive network rerouting (Figure 3I). Overall, these results demonstrate how kinetic modeling can capture the intracellular consequences of targeted therapy and inform the selection of metabolic biomarkers for follow-up studies.

#### Shared Ceramide-linked Stress Response Across Drug Targets and Physiologies

Having established the workflow for the hexokinase inhibition case, we next applied it to TMDS and TRIOK inhibition in both *BRCA1*^WT^ and *BRCA1*^MUT^ model populations (Figures S2–S6). Our goal was to assess whether *BRCA1* status is associated with distinct drug-response patterns and whether shared metabolic shifts occur independently of the specific therapy. To this end, we summarized concentration and flux changes across models using pathway-level enrichment analysis and mapped them to pathways annotated in Recon3D (Figure 4). As anticipated from their distinct biological roles, each target produced characteristic responses. Hexokinase and triokinase inhibition resulted in system-wide shifts in energy and biosynthetic fluxes, while thymidylate synthase inhibition led to nucleotide-specific depletions (Figure 4A, 4B). Interestingly, TRIOK inhibition triggered distinct metabolic rewiring depending on the *BRCA1* physiology: the pentose phosphate pathway was disrupted in *BRCA1*^WT^, while cholesterol and lipid metabolism were significantly affected in *BRCA1*^MUT^ (Figure 4A).

**Figure 4.**
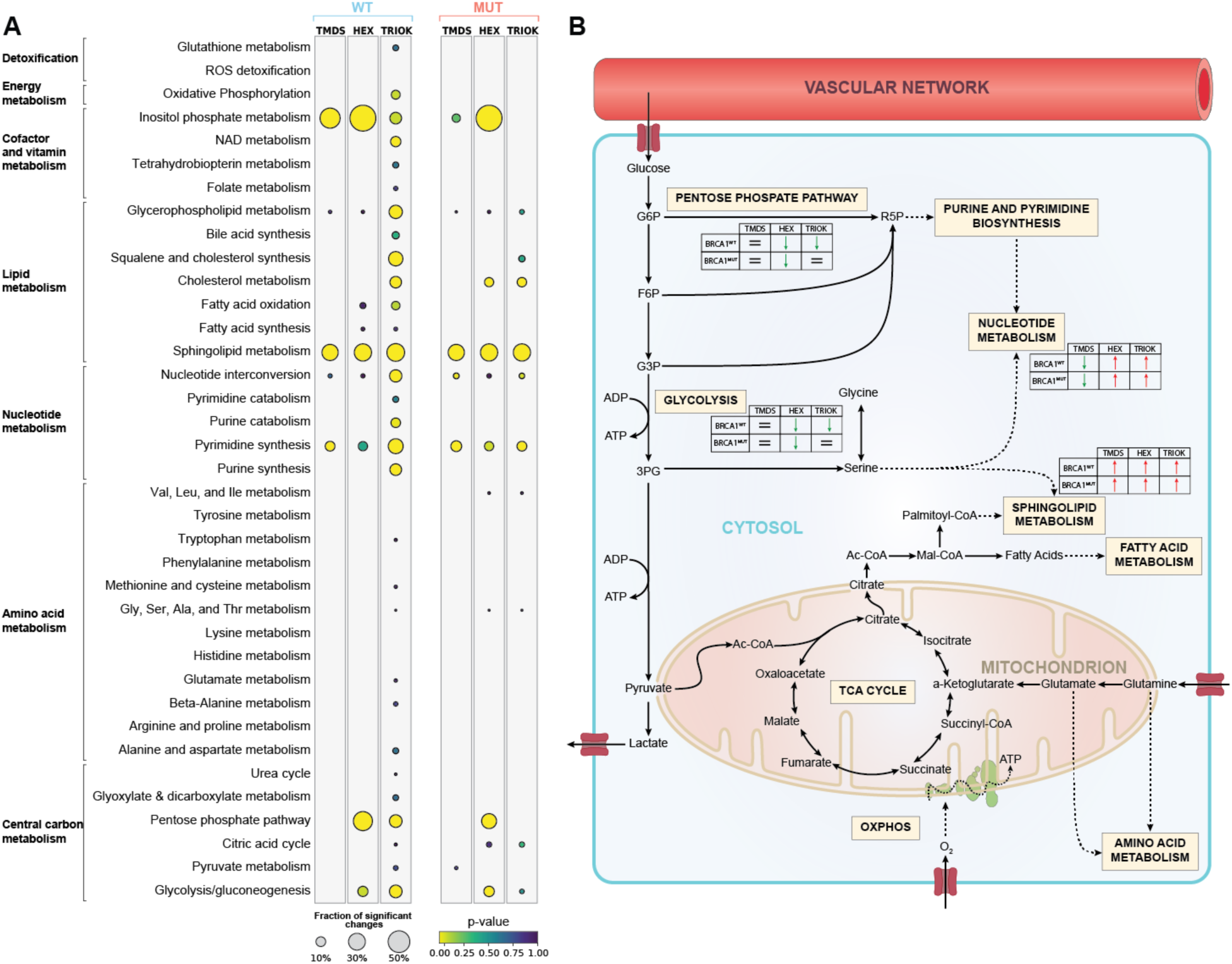
Distinct targets and *BRCA1* physiologies produce characteristic subsystem signatures with a common stress response. (A) Pathway-level enrichment analysis of steady-state metabolite concentration changes predicted across kinetic model populations. Enrichment represents the fraction of consistently deregulated metabolites per pathway (hypergeometric test), with circle size denoting this fraction and color indicating the corresponding p-value. Each drug produces a distinct subsystem signature: for example, TRIOK blockade selectively perturbs the pentose phosphate pathway (PPP) and cholesterol/lipid metabolism in *BRCA1*^MUT^ cells, whereas TMDS inhibition primarily affects nucleotide metabolism in both physiologies. All three interventions consistently deregulate sphingolipid metabolism, reflecting the ceramide-mediated stress response characteristic of diverse chemotherapeutics. (B) Schematic overview of the effects of hexokinases (HEX), triokinase (TRIOK), and thymidylate synthase (TMDS) inhibition on the core metabolic pathways. Red arrows denote increased pathway activity, green arrows denote decreased activity, and equality symbols indicate minimal or no change.

Across all targets and physiologies, sphingolipid metabolism was consistently deregulated, characterized by accumulation of ceramide, dihydroceramide, sphingomyelin, and sphinganine. This pattern aligns with the established role of ceramide in stress responses. Ceramide is a key bioactive lipid that engages signaling pathways governing growth arrest, differentiation, and senescence.^15^ Interestingly, similar to our results, ceramide levels have been found to increase during various drug therapies and across different types of cancer.^16,17,84^ In drug-resistant cancer cells, ceramide is converted into sphingomyelin, which is suggested to help prevent pro-apoptotic signals.^18,85^ While our current models lack the S1P branch, the consistent accumulation of ceramide and its derivatives recapitulates metabolic contexts that mirror experimentally observed drug-induced shifts. Importantly, the model itself does not simulate signaling cascades; however, the models suggest that ceramide accumulation represents a direct metabolic signature of growth arrest, rather than a downstream effect specific to individual drug mechanisms. This shared metabolic buildup provides a mechanistic explanation for why diverse therapies trigger apoptosis through ceramide signaling.

In summary, our population of kinetic models captures how metabolic therapies reshape cancer metabolism, revealing vulnerabilities that depend on cellular context and reproducing known drug-induced shifts in metabolism. By linking dose–response behavior to pathway-level flux and metabolite changes, the models provide mechanistic insights often inaccessible experimentally, offering a robust platform for identifying therapeutic targets, characterizing metabolic biomarkers, and guiding strategies for precision metabolic interventions.

### Cancer phenotype comparisons reveal metabolic and regulatory shifts

*BRCA1* loss-of-function (as in *BRCA1*^MUT^) reshapes ovarian cancer metabolism, manifesting as elevated glycolysis and other malignant traits compared to *BRCA1*^WT^, which retains a less aggressive metabolic state. Yet the extent to which these changes reflect direct regulatory consequences of *BRCA1* loss remains largely unexplored.

To address this, we used the kinetic models to identify the minimal set of enzyme activity changes that can account for the metabolic differences between *BRCA1*^WT^ and *BRCA1*^MUT^ cells. Defining these enzymatic changes provides a mechanistic basis to link altered metabolic states back to upstream regulation by connecting the affected enzymes to transcription factors and ultimately to *BRCA1*.

We quantified metabolic differences by comparing pathway fluxes between *BRCA1*^MUT^ and *BRCA1*^WT^ across 12 major subsystems of central and peripheral metabolism (Figure 5). These pathways, comprising 298 reactions, span commonly studied subnetworks such as glycolysis and mitochondrial energy production^10–12^, but also extend into additional subsystems, providing a more holistic view of the metabolic rewiring that underlies the *BRCA1*^MUT^ phenotype.

**Figure 5.**
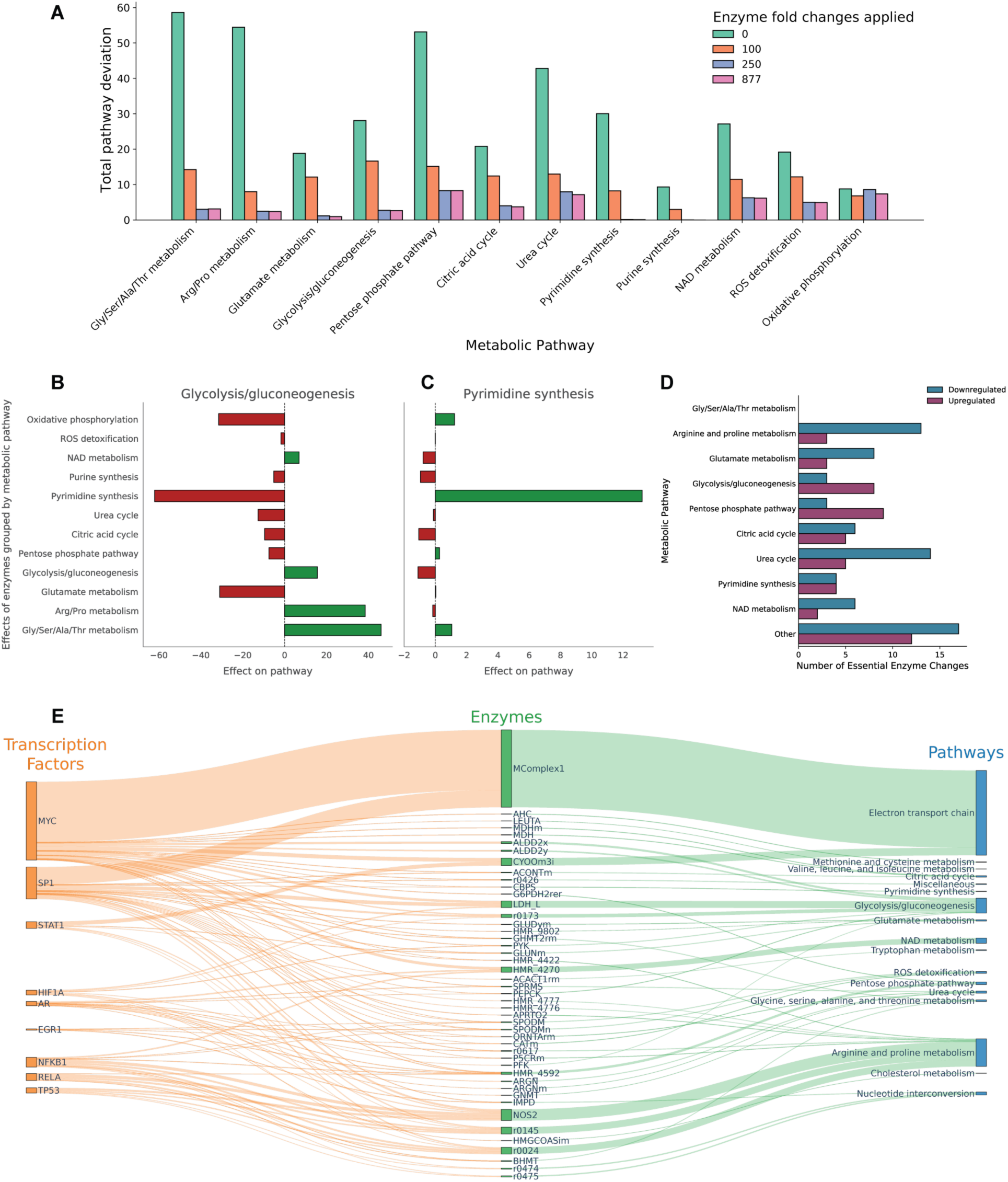
Enzyme level shifts capture the metabolic differences between *BRCA1*^MUT^ and *BRCA1*^WT^, suggesting a *BRCA1*-linked network-wide reprogramming. (A) The residual flux deviation of the 12 core metabolic pathways progressively decreases as more enzyme fold changes are imposed. Allowing 250 enzymes recapitulates most of the *BRCA1*^WT^ flux profile, with little additional gain when all 877 candidate enzymes are utilized, highlighting a limited set of drivers of metabolic shift. (B) Glycolytic fluxes are redirected from the *BRCA1*^MUT^ to the *BRCA1*^WT^ state through coordinated adjustments across neighboring pathways, illustrating distributed network-wide control. (C) Flux deviations in pyrimidine biosynthesis are corrected predominantly by enzymes within the pathway itself, reflecting localized regulatory control. (D) We identified a core set of 145 enzymes that consistently require a fold change. They are grouped by pathway and categorized by up- or downregulation. The predominance of downregulated enzymes is consistent with an overall reduction in metabolic activity in *BRCA1*^WT^ compared to *BRCA1*^MUT^. (E) Sankey diagram illustrates the relationship between the core set of enzymes and their regulating transcription factors (TFs). A subset of *BRCA1*-interacting TFs exerts widespread influence, suggesting that *BRCA1* loss propagates through specific TF hubs to drive network-level metabolic rewiring.

As we allowed progressively more enzymes to change activity, the metabolic state of *BRCA1*^MUT^ became increasingly similar to that of *BRCA1*^WT^ cells (Figure 5A, Figure S9). Strikingly, adjusting 250 enzymes was almost as effective as altering all candidates, capturing most of the differences between the two physiologies (Figure S9). Many pathways showed substantial improvement in flux alignment, and in nucleotide metabolism, specifically purine and pyrimidine biosynthesis, the mutant state could be fully reconciled with the *BRCA1*^WT^ profile (Figure 5A). These findings suggest that a coordinated, yet limited set of enzyme activity shifts is sufficient to account for most of the metabolic reprogramming that distinguishes *BRCA1*^MUT^ from *BRCA1*^WT^ ovarian cancer cells.

The way these flux alignments were achieved depended on the pathway. In glycolysis, alignment with the wild-type state required distributed adjustments across multiple connected pathways, consistent with its role as a central hub in cellular metabolism (Figure 5B). By contrast, pyrimidine biosynthesis was largely brought into alignment through local changes within the pathway itself (Figure 5C). Local regulation is observed in other distal pathways too (Figure S10). This contrast reflects an emergent stoichiometric control pattern: core metabolic pathways are modulated by broad network-level interactions, while terminal pathways are locally regulated. Together, these results suggest that extensive, coordinated adjustments to enzyme activity are required to reconcile the metabolic states of *BRCA1*^MUT^ and *BRCA1*^WT^ cells.

Because metabolism is highly interconnected, there are numerous possible ways to adjust enzyme activity to perturb the *BRCA1* mutant state so that it resembles the wild-type state. Across all possible ways, a consistent group of 145 enzymes always appeared among the top candidates (Figure 5D). These enzymes represent a core set of metabolic regulators that seem essential for understanding how *BRCA1* loss alters ovarian cancer metabolism. Most of them showed lower activity when shifting from the mutant to the wild-type state, which aligns with the observation that *BRCA1* mutant cells grow faster and behave more aggressively, typically relying on higher enzyme activity to meet their demands.^86^ These core enzymes are distributed across multiple pathways (Figure 5D), but amino acid metabolism is especially enriched. This finding is consistent with our broader observation that disturbances in amino acid metabolism account for a significant portion of the metabolic differences between *BRCA1*^MUT^ and *BRCA1*^WT^ cells (Figure 5A).

To link the predicted enzyme-level changes and *BRCA1*-mediated transcriptional regulation, we investigated transcription factors (TFs) known to regulate the expression of the 145 core enzymes. Using the TFLink database^87^, we identified 108 TFs regulating 60 of these enzymes through 182 distinct genes (Data S3). Focusing on the top 20 TFs with the broadest regulatory coverage, we identified six (MYC, STAT1, HIF1A, AR, EGR1, TP53) that are known to interact directly with *BRCA1*. A complementary literature search identified three more *BRCA1*-interacting TFs (SP1, RelA, and NFKB1) that engage *BRCA1* through protein-protein interactions. Specifically, *BRCA1* physically interacts with SP1, which interferes with SP1’s ability to transactivate IGF-IR.^88^ This may have novel downstream effects stemming from *BRCA1* action. *BRCA1* has also been shown to bind the p65/RelA subunit of NF-κB^89^ and act as a transcriptional co-activator. Interestingly, it has been demonstrated that the absence of functional *BRCA1* leads to an increase in the expression of NF-κB target genes.^89,90^

Collectively, these nine *BRCA1*-associated TFs regulate 46 of the 145 core enzymes, suggesting a possible transcriptional mechanism through which *BRCA1* could exert control over a significant fraction of the metabolic network (Figure 5E). Notably, MYC and STAT1 could mediate strong effects on the electron transport chain, while glycolysis and several amino acid biosynthesis pathways are also prominently regulated by this TF set.

Although *BRCA1*-linked transcription factors explained only part of the predicted enzyme activity changes, this is not unexpected, given the gaps in current interaction maps, the missing gene annotations for 24 of the 145 enzymes, and indirect influences such as feedback loops or post-transcriptional regulation. Together, these findings point to a broader and more complex regulatory network that connects *BRCA1* to ovarian cancer metabolism. Importantly, the modeling condenses this complexity into a clear set of enzyme activity shifts that can now be tested experimentally. Approaches such as ^13^C-tracing, targeted proteomics, or CRISPR-based transcription factor screens provide a practical path to uncover how *BRCA1* loss drives regulatory and metabolic reprogramming in ovarian cancer.

## Discussion

Large-scale kinetic modeling constrained by multi-omics and physicochemical data offers a transformative shift in how we study metabolism by enabling near-genome-scale, time-resolved simulation of fluxes and metabolite concentrations. Populations of physiologically consistent kinetic models captured the observed heterogeneity of the two isogenic cell lines, linking flux and metabolite concentration dynamics. Using Metabolic Control Analysis (MCA), we identified 28 enzymes exerting the strongest influence over proliferation; 24 of these correspond to market-approved or experimental drugs, validating the framework, while the remaining four represent unexplored metabolic vulnerabilities. Compared to gene knockout simulations in steady-state models, which identified hundreds of potential targets (417 for *BRCA1*^WT^ and 416 for *BRCA1^MUT^*, Data S4), MCA identified a smaller, mechanistically interpretable set of enzymes with strong control over proliferation. These results highlight the enhanced resolution and mechanistic depth of kinetic modeling, which captures nonlinear and regulatory effects that are inaccessible to stoichiometric models.

Dynamic modelling reveals emergent metabolic signatures of therapeutic stress, accurately reproducing experimentally observed responses while providing mechanistic insight into how therapies reshape cancer-cell metabolism. Importantly, these insights may generalize beyond ovarian cancer to other proliferative or metabolic disease contexts. By tracking all intracellular fluxes and metabolite levels, the models provide a mechanistically detailed, systems-wide view of how drug therapies impact the entire metabolic network, revealing compensatory adaptations such as increased mitochondrial respiration following hexokinase inhibition and ceramide-centered sphingolipid accumulation associated with stress responses. Collectively, these simulations demonstrate that large-scale kinetic modelling can both validate known treatment effects and uncover previously uncharacterized network-level metabolic responses, offering a robust platform for hypothesis generation and therapeutic exploration.

Finally, a minimal set of enzyme fold changes explained the differences in the metabolic states between *BRCA1*^MUT^ and *BRCA1*^WT^ physiologies, and we proposed a core set of TFs that are potentially mediated by *BRCA1* to achieve the observed metabolic shifts. This allows us to focus on a much narrower set of TFs, especially because *BRCA1* interacts with dozens of transcription factors (TFs) that control hundreds of metabolic genes. According to TFLink^87^, *BRCA1*-associated TFs regulate 872 Recon3D genes encoding more than 2,000 enzymes. However, such associations alone cannot reveal which enzyme activity changes are responsible for the distinct metabolic behavior of *BRCA1*^MUT^ and *BRCA1*^WT^ cells. This framework provides a blueprint for dissecting how transcriptional-regulatory networks impose dynamic changes on metabolic networks, bridging gene regulation with metabolite and flux dynamics in a unified view, and can be extended to study enzymatic rewiring in other contexts, including metabolic disease and tissue adaptation.

Overall, these results establish kinetic modeling as a dynamic, mechanistic complement to conventional genome-scale models, providing a resource for hypothesis generation, target prioritization, and pharmacological exploration across metabolic disease contexts.

This study has several limitations that should be considered when interpreting the results. First, the current models are based on multi-omics data from an isogenic pair of ovarian cancer cell lines (*BRCA1*^WT^ and *BRCA1*^MUT^). This design enables isolation of the effects of *BRCA1* loss but restricts generalizability, as broader validation with patient-derived samples and other tumor contexts may be needed to extend the predictions. Second, although the models incorporate detailed mechanistic rate laws and are constrained by multi-omics and physicochemical data, they do not explicitly account for enzyme regulation through allosteric interactions or signaling cascades, both key components in mammalian physiology. Nonetheless, despite not including these layers of regulation, the models reliably reproduced hallmark drug responses and physiological phenotypes, indicating that much of the observed system behavior can be explained by enzyme abundances and metabolic network structure alone. Finally, the models quantify *BRCA1*-dependent differences in ovarian cancer metabolism across nearly 300 reactions spanning 12 subsystems. In contrast, most prior studies have focused on glycolytic and oxidative pathways, often describing *BRCA1*-associated shifts simply as enhanced glycolysis. This difference may complicate direct comparison with earlier reports, but it provides an unprecedented mechanistic resolution and generates experimentally testable hypotheses for future work.

### Future and implications

Kinetic models provide a platform for generating mechanistic predictions that guide experimental and clinical investigations. Incorporating explicit pharmacokinetic and pharmacodynamic (PK/PD) modules, such as expressions that track drug absorption, distribution, metabolism, and clearance, will enhance the relevance of the predictions, facilitate dose optimization, and support the time-consuming and expensive drug development process. Integrating patient-specific data could enable personalized models, tailored to each patient’s physiology, linking predicted metabolic vulnerabilities to customized drug therapies. Efforts are underway to accelerate model construction and parameterization using machine learning^91–93^, streamlining the workflow from data acquisition to actionable predictions.

Cancer metabolism is shaped by oncogenic signaling, which regulates pathway activity, proliferation, and flux distributions. Key signaling pathways frequently activated in tumors, such as the PI3K–AKT–mTORC1 and MYC pathways, increase metabolic flux through numerous pathways to support cell proliferation.^4,5^ Metabolites themselves can feed back onto signaling cascades, thereby creating an interconnected network that facilitates tumor progression.^4,5^ Despite this, current models remain mostly small-scale or cascade-specific, highlighting the need for a robust workflow to develop dynamic, large-scale kinetic models that integrate signaling and metabolism.^94^

Kinetic models also enables exploration of how the dynamic availability of nutrients shapes cancer metabolism. Cancer cells utilize various carbon sources, including glucose, lactate, amino acids, and proteins, to sustain proliferation.^5,64^ Modeling how fluxes adjust to shifts in primary carbon sources or dietary regimes could reveal previously unrecognized metabolic vulnerabilities and inform the development of novel pharmacological interventions.^95,96^

Simulating interactions between cancer cells and the tumor microenvironment (TME) could further clarify context-dependent metabolic adaptations. Numerous, yet incompletely understood, mechanisms enable cancer cells to exploit their neighboring cells to increase their chances of survival.^45,97,98^ Cancer cells compete with immune cells, such as activated T cells, for key nutrients such as glucose and amino acids.^45^ They also co-opt the metabolism of neighboring stromal cells, which secrete numerous metabolites, including lactate and glutamine.^45^ Incorporating the metabolism of these primary TME cell types into future kinetic models would open new opportunities to design therapies that target tumor metabolism without compromising immune function.^46,53^

### Conclusions

Near-genome-scale kinetic models constrained by tissue- or patient-specific multi-omics data extend conventional genome-scale frameworks by capturing the dynamic behavior of metabolic networks. The ovarian cancer models presented here, validated against diverse experimental and clinical observations, provide a robust mechanistic platform for interrogation, hypothesis generation, and rational therapeutic design. They hold strong potential for advancing precision oncology and personalized medicine by enabling network-level, dynamic insights into cancer metabolism and illustrating how dynamic metabolic modelling can guide interventions across metabolic diseases and tissue physiology more broadly.

## Materials and Methods

### Reduced metabolic network and multi-omics data integration

The first step in our ORACLE workflow^99^ for constructing kinetic models involves defining the reduced metabolic network that contains the pathways of interest for a given biological question. The next step is the integration of available omics data by applying methods adapted for integrating transcriptomics (REMI^40^, iMAT^100,37,101^, INIT^102^, ΔFBA^103^), metabolomics (TFA^104^, INTEGRATE^105^), and fluxomics^106^ data. In this study, we employed a reduced stoichiometric model of ovarian cancer metabolism integrating omics data as previously described.^56^ Starting from the Recon3D^57^ genome-scale model of human metabolism, those authors applied the redGEM^107^, lumpGEM^108^, and redGEMX^54^ tools to generate a reduced network tailored to ovarian cancer. They then integrated transcriptomic data from *BRCA1*^WT^ and *BRCA1*^MUT^ ovarian cancer cell lines using the iMSEA workflow^56^, which consistently merges transcriptomics, metabolomics, and fluxomics information through TFA-^104^ and REMI^40^-based constraints. The resulting stoichiometric model comprised 2,369 reactions and 974 metabolites distributed across eight cellular compartments and 64 metabolic subsystems.

In the present study, we further integrated experimentally measured metabolite levels of ATP, ADP, and AMP from Park et al.^109^ We additionally imposed a basal nonzero flux constraint (10^-6^ mmol·gDW⁻¹·h⁻¹) for all reactions to enforce minimal enzyme activity and ensure global network connectivity. As a result, some reactions lost their ability to carry flux and were removed from the models, yielding final network sizes of 2,359 reactions for the *BRCA1*^WT^ physiology and 2,344 for the *BRCA1*^MUT^ physiology.

### Determining reference metabolic steady states

Many intracellular reactions can operate in both directions. Even after integrating multi-omics and physicochemical data along with expert knowledge, a significant number of reactions remained bidirectional (BDRs), resulting in considerable flux uncertainty. To reduce the feasible solution space, we applied an iterative, criteria-driven procedure to assign reaction directionalities:

1. Exchange-related reactions. For BDRs connecting intracellular metabolites to the extracellular medium, directionality was set based on the growth medium: metabolites absent from the medium were assumed to be secreted.
2. Futile-cycle elimination. We identified reaction pairs whose opposing directions could result in artificial ATP dissipation, e.g., ATP-driven import opposed by passive export. When such futile cycles were identified, we aligned their directions to eliminate them and maintain energetic consistency.
3. Flexibility-based selection. For the remaining BDRs, we performed flux variability analysis (FVA) and, wherever possible, assigned the direction with the larger feasible range, thereby favoring greater flexibility.

It is important to note that assigning a directionality to a reaction does not restrict it to operate exclusively in that direction. The specified directionality defines only the reference orientation of fluxes at the sampled steady state. For reactions governed by reversible kinetic rate laws (e.g., Reversible Michaelis-Menten, Convenience kinetics^110^), the net flux can reverse when the substrate-to-product concentration ratio changes sufficiently. Thus, although a reference direction is chosen for computing the steady-state solution, reaction directionalities remain dynamically adaptable to environmental or pharmacological perturbations.

Once the reference reaction directionalities were established, we sampled steady states consistent with the integrated experimental constraints, thereby reproducing the observed metabolic heterogeneity. To achieve this, we utilized the Artificial Centering Hit-and-Run (ACHR) sampler implemented in the pyTFA toolbox^39^ to generate 5,000 steady-state samples with a thinning step of 500. Each sample included flux and metabolite concentration values across the network, as well as thermodynamic information such as ΔG and ΔG°.

### Generation of the kinetic model scaffold

The reduced metabolic network served as the scaffold for building the kinetic model. Each reaction was assigned an appropriate rate law mechanism from a comprehensive library implemented in the SKiMpy toolbox.^60^ By default, SKiMpy assigns the enzyme mechanism based on the reaction’s stoichiometry. The produced kinetic model consists of 716 mass balances (and 25 pool-conservation equations for dependent metabolites) governed by 2,131 rate laws. In total, the model required values for 9,594 kinetic parameters, including 7,463 Michaelis-Menten constants (K_m_), 2,131 maximum velocity constants (v_max_), and 2,193 thermodynamic equilibrium constants (K_eq_).

### Generation of a population of kinetic models

To generate kinetic parameter values, we applied the ORACLE workflow.^99^ Using a steady-state sample of fluxes and concentrations, we generated 100 independent parameter sets, each of which was consistent with the imposed steady state. To evaluate their physiological relevance, we analyzed the local dynamics of each model by computing the eigenvalues of the system’s Jacobian. We retained models based on physiologically relevant dynamics: models were considered physiologically relevant if their dominant time constant (*τ*_*d*_) was at least three times faster than the doubling time of the modeled cancer cell.^92,93,111,112^

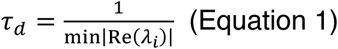

Because some steady-state samples did not yield physiologically relevant kinetic models, we repeated this process for each physiology separately until the desired distributions were obtained, as described in the next section. The final number of kinetic models was 13,731 for the *BRCA1*^WT^ physiology, derived from 556 steady-state samples, and 6,732 for the *BRCA1*^MUT^ physiology, derived from 396 steady-state samples.

### Metabolic Control Analysis

Metabolic Control Analysis (MCA) was performed using the SKiMpy toolbox to quantify how changes in enzyme activity propagate throughout the network. The sensitivity of fluxes and metabolite concentrations to enzyme perturbations is expressed through the control coefficients:^62,113^

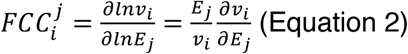

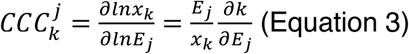

where *FCC_i_^j^*^-^ and *CCC_i_^j^*^-^ denote the flux and concentration control coefficient, respectively. 𝑣*_i_* is the flux of reaction *i*, *E_j_* the enzyme concentration *j*, and *x_k_* is the concentration of metabolite *k* at the reference steady state. A positive control coefficient indicates that the flux or concentration is expected to increase with an increase in enzyme concentration. For example, the flux control coefficient of the growth rate with respect to triokinase (TRIOK) for the *BRCA1*^WT^ physiology is 0.72 (Figure 1A), meaning that a 2-fold increase in TRIOK concentration would result in an approximately 1.65 times increase in the growth rate relative to the reference state.

MCA predictions are accurate only for small perturbations around the reference state. Nevertheless, the magnitude and sign of the control coefficients provide valuable insight into enzymes that affect cancer proliferation. The actual impact of larger perturbations was further assessed by performing drug simulations. Different steady-state samples and kinetic parameter values will yield different values for the FCCs and CCCs. In the analysis, we report the population averages and the interquartile ranges.

To evaluate the effect of top enzymes on the biomass precursors, we calculated their CCCs (Eq. 3) and the control coefficients of their synthesis rate:

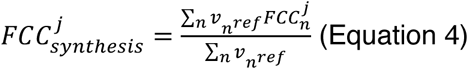

where n denotes the number of reactions that are producing the biomass building block, and the 𝑣*_n_^ref^* is the flux value at the reference state.

For every kinetic model that satisfied the local dynamic criteria, we calculated all the flux and concentration control coefficients across the network. To identify statistically significant trends, we evaluated the convergence of the distributions of the flux control coefficients of growth rate with respect to each enzyme. To ensure robustness, outlier control coefficient values were removed using the interquartile range (IQR) method, applied independently to each enzyme’s flux control coefficient distribution. Convergence was considered achieved when the first three moments (mean, variance, skewness) of each distribution changed by less than 10% upon addition of a new kinetic model to the population. This allowed us to discuss the averages and the standard deviations of the key enzymes for cancer growth, which were picked based on the absolute average values of the control coefficients. For these key enzymes, we further calculated the average flux and concentration control coefficients for all network variables (fluxes or concentrations), allowing us to obtain a system-level understanding of their metabolic effects.

### DrugBank data extraction

For each of the 28 drug targets predicted by the MCA workflow, we retrieved associated drugs based on information from DrugBank (version 5.1.13, released on January 2, 2025).^65^ The 28 enzymes were mapped to their Uniprot identifiers, and checked which drugs are associated with them. For each drug, we also extracted its ATC code, approval status (approved or experimental), and documented mode of action, whenever available.

### *In silico* drug administration simulations

We performed the drug administration simulations on a representative subset of the original kinetic model population, selected through a stratified sampling approach. Analogous to clinical studies, where participants are recruited to capture variability across defined strata, we divided the models according to their initial glycolytic and mitochondrial ATP yields. Glycolytic ATP yield is calculated as the ratio of the sum of PYK and PGK fluxes to the glucose uptake rate. Similarly, the mitochondrial ATP yield was calculated as the ratio of the mitochondrial ATP synthase (ATPS4mi) flux to the glucose uptake rate. We then divided the observed ranges into three strata for each yield, resulting in nine possible combinations, and randomly selected 50 steady-state samples in total, spread across the nine metabolic subtypes. Importantly, because the number of steady-state samples varied across strata, we ensured that the 50 representative metabolic states reflected the relative density of models within each stratum. This resulted in 972 kinetic models for the *BRCA1*^WT^ physiology and 1,134 for the *BRCA1*^MUT^ physiology.

To simulate the effects of anticancer drugs, we assumed that the drug action directly impacts the enzymatic activity of the target enzyme. This effect was modeled by introducing an additional ODE that exponentially reduces the enzyme concentration toward a specified target value:

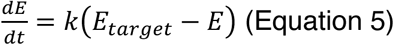

where *k* is the rate constant of the effect, and *E*_*target*_ is the target enzyme concentration achieved due to the drug action. Since the models are formulated in terms of V_max_ values rather than explicit enzyme concentrations, we assumed an initial enzyme concentration of 1 M. Under this assumption, the catalytic rate constant, k_cat_, is equal to the sampled V_max_ value. We subsequently integrated all kinetic models using as initial conditions the steady state sample around which each model was built. Numerical integration was performed using CVODE^114^ over a simulation time of 25 model days, providing ample time for the system to reach a new steady state. In practice, most models settled into a new metabolic state within one doubling time. This procedure yielded the time evolution of all the concentrations and fluxes in the metabolic network across the subpopulation of kinetic models.

To estimate EC_50_ values for each drug, we performed drug administration simulations by progressively decreasing *E*_*target*_ from 0.999 (effectively no perturbation) to 10^-9^ (complete inhibition). Cell viability was calculated as the ratio of the initial to the final growth rate. The resulting data points were used to fit a 4-parameter logistic model, following the format described by Corsello et al.^81^

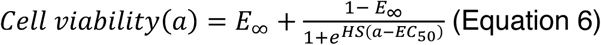

where *a* denotes the target enzyme concentration (*E*_*target*_), *E*_∞_ the residual effect when the enzyme is completely inhibited, *EC*_50_ the final enzyme concentration that produces a half-maximal effect (the cell viability will be halfway between 1 and *E*_∞_), and *HS* is the hill slope that determines the steepness of the dose-response curves.

This analysis across all kinetic models yielded a distribution of pharmacodynamic parameters, capturing the expected ranges of drug-induced growth inhibition. The reported values for each parameter represent the average and standard deviation after removing outliers, which were excluded using the interquartile range method.

### Volcano plots

To assess the system-level effects of drug target inhibition, we created volcano plots that show the significantly perturbed fluxes and concentrations within the network. For each model, we calculated log_2_ fold changes in metabolite concentrations and log_10_ fold changes in reaction fluxes between the initial and final metabolic states. A one-sample t-test was applied across all the models to identify significantly perturbed concentrations and fluxes, and p-values were corrected for multiple testing with the Benjamini–Hochberg method.^115^ To visualize the distance of each metabolite and flux from the target enzyme, we utilized the adjacency matrices generated by the redGEM workflow^107^ and calculated the minimum distance between a reaction or metabolite from the metabolites participating in the inhibited reaction.

### Pathway enrichment analysis

To compile and compare the changes in metabolic pathways upon drug treatment, we conducted pathway enrichment analysis. Metabolites with an absolute log_2_ concentration fold change ≥ 2 and q ≤ 0.01 were considered significantly perturbed and were mapped to pathways through the Recon3D GEM. We calculated the fraction of significantly altered metabolites within each metabolic pathway and used a hypergeometric statistical test to determine whether the number of perturbed metabolites exceeded what would be expected by chance. To determine whether each pathway was predominantly up- or downregulated following treatment, we examined both the direction and magnitude of metabolite fold changes within each pathway. In nearly all cases, the majority of metabolites and fluxes within a pathway changed in the same direction, allowing us to assign an overall qualitative trend of up- or downregulation. These summarized effects were then visualized on a simplified metabolic map, providing an intuitive overview of how drug treatment reshaped intracellular metabolism.

### Comparison of *BRCA1*^WT^ and *BRCA1*^MUT^ phenotypes

To identify enzyme fold changes that will minimize flux deviations between *BRCA1*^WT^ and *BRCA1*^MUT^ phenotypes, we used the NRA formulation^116,117^ implemented in the NOMAD framework.^111^ The NRA requires as inputs the reference *BRCA1*^MUT^ flux state (serving as the initial metabolic state) and a kinetic parameter set (used to calculate the control coefficients of the network), which together enable the estimation of the new metabolic state after applying specified enzyme fold changes. For comparison, the target flux state was defined from *BRCA1*^WT^ physiology.

We calculated the flux deviations for each pair of *BRCA1*^WT^ and *BRCA1*^MUT^ steady states that were sampled for the MCA workflow, resulting in a distribution of deviation values. From this distribution, we selected the representative pair that was closest to the average of the distribution.

We then selected a physiologically relevant kinetic parameter set and used it to construct a kinetic model centered on the selected *BRCA1*^MUT^ steady-state sample. A series of robustness tests was performed to determine whether the chosen kinetic parameter set was suitable for the subsequent analysis. Specifically, we randomly perturbed each metabolite concentration by up to ±10% and simulated the system’s response. A response was considered robust if the model returned to its original flux and concentration profile within one doubling time. This procedure was performed 100 times, and in all cases the model was able to return to its original steady state. Having passed this test, the kinetic model was deemed robust and was subsequently used in the NRA formulation.

NRA was used to predict the new flux distribution of the metabolic network following perturbations in enzyme activity. For this purpose, we optimized the fluxes of 298 reactions belonging to the 12 core pathways, representing key features of cancer metabolism activity. We allowed 877 enzymes to vary in activity, comprising all enzymes catalyzing biotransformations in our reduced model and 112 enzymes catalyzing the transport of metabolic species, and whose genes exhibited significant expression fold changes between the ovarian cancer cell lines. Enzyme upregulation was constrained to a maximum 50-fold increase, based on the maximum gene expression fold change observed in our reference gene expression data, while downregulation was allowed to effectively knock out the enzyme. To guide NRA toward a solution producing a flux distribution closest to the *BRCA1*^WT^ physiology, we used the following objective function:

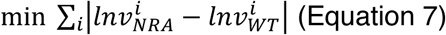

where 𝑣_*NRA*_ is the new flux solution produced by NRA, and 𝑣_*WT*_ is the reference *BRCA1*^WT^ flux profile. The fluxes are represented on a log scale, as NRA operates in a log-linear space. We introduced additional constraints to limit the number of enzymes allowed to have a fold change, enabling the identification of the minimal set of perturbations required to minimize the distance between the *BRCA1*^WT^ and *BRCA1*^MUT^ flux distributions. This analysis revealed that applying a fold change to 250 of the 877 enzymes produced results nearly indistinguishable from those obtained when all enzymes were permitted to vary, indicating that a subset of enzymes exerts dominant control over the phenotypic differences between the two physiologies.

To quantify the contribution of NRA-predicted enzyme fold changes to pathway-level alignment between physiologies, we defined pathway effect scores. For a given target pathway, the effect from a source (contributing) pathway was calculated as:

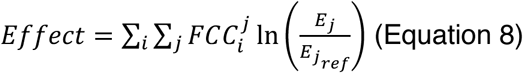

with *i* indexing all fluxes in the target pathway and *j* indexing all enzymes in the source pathway. Positive effect scores indicate that enzymes in the source pathway collectively push upward fluxes in the target pathway, whereas negative scores indicate a downward influence. Central pathways, such as glycolysis, typically integrate contributions by multiple source pathways, whereas biosynthetic pathways, like pyrimidine synthesis, are predominantly driven by enzymes within the same pathway.

### Identifying a minimal core set of enzyme fold changes

Due to the strong interconnectivity of the metabolic network, multiple combinations of 250 perturbed enzymes could reproduce similar alignment between the two phenotypes. To identify the enzymes that were consistently required, we developed an iterative workflow. Starting with one solution set of 250 enzymes, we systematically blocked the fold change of each enzyme and re-optimized for Equation 7 using the NRA formulation. If the new solution achieved an objective value within 5% of the original, the removed enzyme was considered non-essential for reproducing the flux alignment. Conversely, if removing the enzyme prevented the optimization from meeting this criterion, the enzyme was deemed necessary and considered active across all alternative solutions. This procedure identified a robust core of 145 enzymes that consistently appeared across solutions, highlighting them as key determinants of how BRCA1 loss reshapes ovarian cancer metabolism.

### TFLink data extraction

We used the TFLink database^87^ to retrieve information on transcription factor (TF) interactions with the metabolic genes of the network, as well as with *BRCA1*. We used information flagged as *small-scale evidence* to ensure that the interactions reported are well-characterized.

## Acknowledgements

This work was supported by funding from the Swiss National Science Foundation grants 200021_188623 and CRSII5_198543, the European Union’s Horizon 2020 research and innovation programme under grant agreement 814408, and the Ecole Polytechnique Fédérale de Lausanne (EPFL).

## Data and Code availability

The data supporting this study’s findings are publicly available in the Zenodo repository at https://zenodo.org/records/17304777 and the links therein.

The code needed to reproduce this study’s results is publicly available at https://github.com/EPFL-LCSB/human-cancer-kinetic-models. The models were built and analyzed using the SKiMpy toolbox, available at https://github.com/EPFL-LCSB/skimpy. NRA is implemented within the NOMAD workflow, which can be accessed at https://github.com/EPFL-LCSB/NOMAD.

## Abbreviations

All metabolite, reaction, and enzyme abbreviations used in this study are consistent with those defined in the human genome-scale reconstruction (Recon3D) and are retained in the curated kinetic models. The full list of abbreviations can be accessed directly within the model files provided with this publication. The same standardized abbreviations are also available through the BiGG/VMH database (https://vmh.life/#home).

## Contributions

Conceptualization and methodology, I.T., M.M., V.H., and L.M.; software, I.T.; data curation, I.T.; data analysis, I.T., M.M., V.H., and L.M.; visualization, I.T. and L.M.; writing — original draft, I.T.; writing — review & editing, I.T., M.M., V.H., and L.M.; supervision, M.M., V.H., and L.M.; funding acquisition, V.H., and L.M.

## Declaration of generative AI and AI-assisted technologies in the writing process

During the preparation of this work, the authors used ChatGPT and Grammarly to improve the readability and language of the manuscript. After using this tool, the authors reviewed and edited the content as needed and take full responsibility for the content of the published article.

## Supplemental information

**Document S1. Figures S1-S10 and Tables S1-S2**

**Data S1. Predicted metabolic drug targets and their associated drugs, as cataloged in DrugBank**

**Data S2. Annotated list of antimetabolites cataloged in DrugBank that were not predicted by the models**

**Data S3. Core set of enzyme fold changes predicted by the NRA formulation, and their connections to transcription factors and BRCA1**

**Data S4. Essential genes identified for both physiologies**

