## Supplemental Figures S1-S10 and Tables S1-S2 for "Multi-omics-driven kinetic modeling reveals metabolic adaptations and vulnerabilities in BRCA1-deficient ovarian cancer" (from another version of this preprint: version 1, 2025-10-23)

### Supplemental information

Multi-omics–driven kinetic modeling reveals metabolic vulnerabilities and differential drug-response dynamics in ovarian cancer

Ilias Toumpe^1^, Maria Masid^2,3^, Vassily Hatzimanikatis^1,3^, Ljubisa Miskovic^1,3^

^1^Laboratory of Computational Systems Biotechnology, École Polytechnique Fédérale de Lausanne (EPFL), Lausanne, Switzerland

^2^Ludwig Institute for Cancer Research, Department of Oncology, University of Lausanne and Lausanne University Teaching Hospital (CHUV), Lausanne, Switzerland

^3^Corresponding authors. Correspondence:

#### Supplementary figures

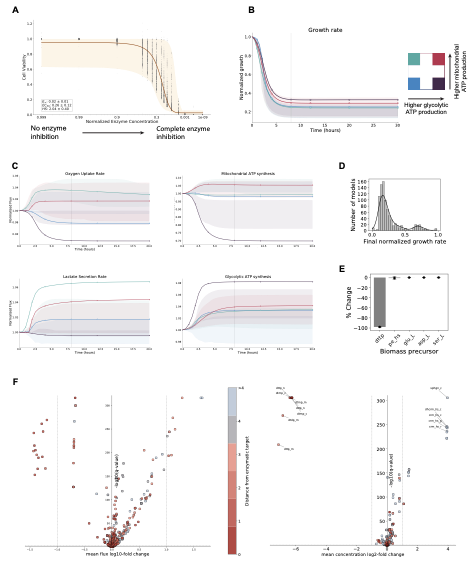

**Figure S1. Thymidylate synthase inhibition in *BRCA1*^WT^ ovarian cancer models.** (A) Dose–response curves of cell viability from 972 kinetic models, plotted against residual TMDS activity. Grey dots represent individual model responses; orange-shaded ranges denote fitted logistic curves, with the average curve highlighted. (B) Transient growth trajectories following a 10-fold reduction in enzyme activity, stratified by glycolytic vs. mitochondrial ATP production initial states (colored boxes). Shaded areas indicate interquartile ranges, and solid lines trace the model closest to each group’s average. Transient responses of (C) oxygen consumption rate (OCR), (D) mitochondrial ATP synthesis, (E) lactate secretion rate, and (F) glycolytic ATP synthesis following the same inhibition. (G) Final growth rate distribution across the population under 10-fold inhibition. (H) Changes in biomass precursor pools following inhibition. Bars denote the average percent change, and the error ranges indicate the interquartile range for the five most affected precursors. (I-J) Volcano plots of steady-state changes in (I) reaction fluxes and (J) metabolite concentrations. Dots are color-coded by network distance from the inhibited hexokinases.

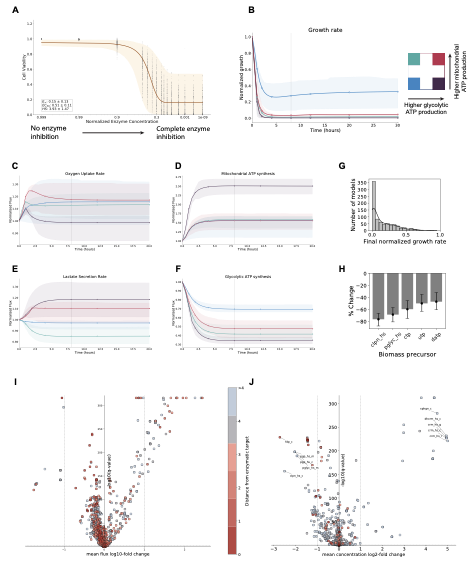

**Figure S2. Triokinase inhibition in *BRCA1*^WT^ ovarian cancer models.** (A) Dose–response curves of cell viability from 972 kinetic models, plotted against residual TRIOK activity. Grey dots represent individual model responses; orange-shaded ranges denote fitted logistic curves, with the average curve highlighted. (B) Transient growth trajectories following a 10-fold reduction in enzyme activity, stratified by glycolytic vs. mitochondrial ATP production initial states (colored boxes). Shaded areas indicate interquartile ranges, and solid lines trace the model closest to each group’s average. Transient responses of (C) oxygen consumption rate (OCR), (D) mitochondrial ATP synthesis, (E) lactate secretion rate, and (F) glycolytic ATP synthesis following the same inhibition. (G) Final growth rate distribution across the population under 10-fold inhibition. (H) Changes in biomass precursor pools following inhibition. Bars denote the average percent change, and the error ranges indicate the interquartile range for the five most affected precursors. (I-J) Volcano plots of steady-state changes in (I) reaction fluxes and (J) metabolite concentrations. Dots are color-coded by network distance from the inhibited hexokinases.

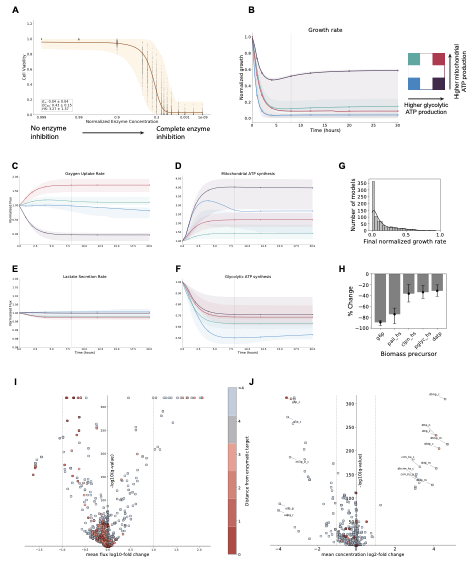

**Figure S3. Hexokinase inhibition in *BRCA1*^MUT^ ovarian cancer models.** (A) Dose–response curves of cell viability from 1,134 kinetic models, plotted against residual hexokinase activity. Grey dots represent individual model responses; orange-shaded ranges denote fitted logistic curves, with the average curve highlighted. (B) Transient growth trajectories following a 10-fold reduction in enzyme activity, stratified by glycolytic vs. mitochondrial ATP production initial states (colored boxes). Shaded areas indicate interquartile ranges, and solid lines trace the model closest to each group’s average. Transient responses of (C) oxygen consumption rate (OCR), (D) mitochondrial ATP synthesis, (E) lactate secretion rate, and (F) glycolytic ATP synthesis following the same inhibition. (G) Final growth rate distribution across the population under 10-fold inhibition. (H) Changes in biomass precursor pools following inhibition. Bars denote the average percent change, and the error ranges indicate the interquartile range for the five most affected precursors. (I-J) Volcano plots of steady-state changes in (I) reaction fluxes and (J) metabolite concentrations. Dots are color-coded by network distance from the inhibited hexokinases.

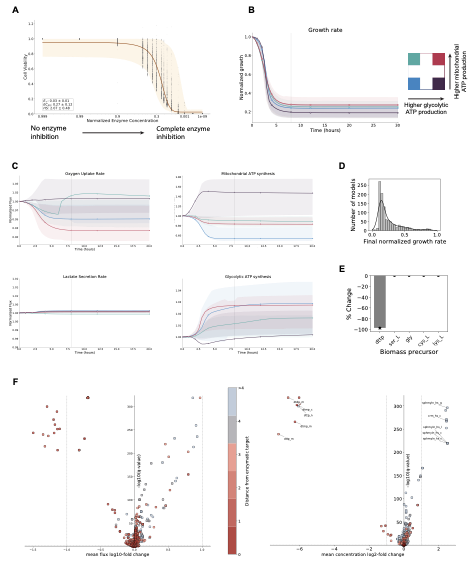

**Figure S4. Thymidylate synthase inhibition in *BRCA1*^MUT^ ovarian cancer models.** (A) Dose–response curves of cell viability from 1,134 kinetic models, plotted against residual TMDS activity. Grey dots represent individual model responses; orange-shaded ranges denote fitted logistic curves, with the average curve highlighted. (B) Transient growth trajectories following a 10-fold reduction in enzyme activity, stratified by glycolytic vs. mitochondrial ATP production initial states (colored boxes). Shaded areas indicate interquartile ranges, and solid lines trace the model closest to each group’s average. Transient responses of (C) oxygen consumption rate (OCR), (D) mitochondrial ATP synthesis, (E) lactate secretion rate, and (F) glycolytic ATP synthesis following the same inhibition. (G) Final growth rate distribution across the population under 10-fold inhibition. (H) Changes in biomass precursor pools following inhibition. Bars denote the average percent change, and the error ranges indicate the interquartile range for the five most affected precursors. (I-J) Volcano plots of steady-state changes in (I) reaction fluxes and (J) metabolite concentrations. Dots are color-coded by network distance from the inhibited hexokinases.

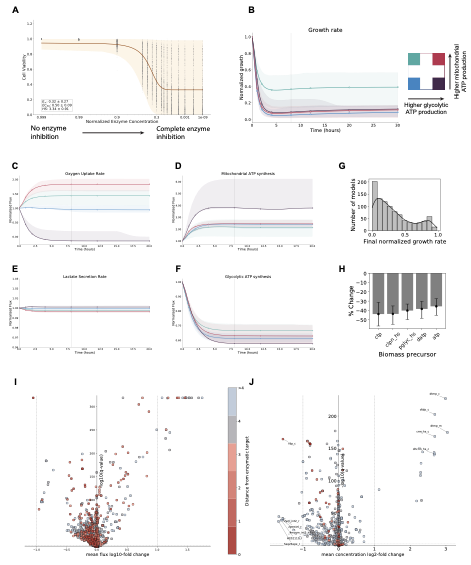

**Figure S5. Triokinase inhibition in *BRCA1*^MUT^ ovarian cancer models.** (A) Dose–response curves of cell viability from 1,134 kinetic models, plotted against residual TRIOK activity. Grey dots represent individual model responses; orange-shaded ranges denote fitted logistic curves, with the average curve highlighted. (B) Transient growth trajectories following a 10-fold reduction in enzyme activity, stratified by glycolytic vs. mitochondrial ATP production initial states (colored boxes). Shaded areas indicate interquartile ranges, and solid lines trace the model closest to each group’s average. Transient responses of (C) oxygen consumption rate (OCR), (D) mitochondrial ATP synthesis, (E) lactate secretion rate, and (F) glycolytic ATP synthesis following the same inhibition. (G) Final growth rate distribution across the population under 10-fold inhibition. (H) Changes in biomass precursor pools following inhibition. Bars denote the average percent change, and the error ranges indicate the interquartile range for the five most affected precursors. (I-J) Volcano plots of steady-state changes in (I) reaction fluxes and (J) metabolite concentrations. Dots are color-coded by network distance from the inhibited hexokinases.

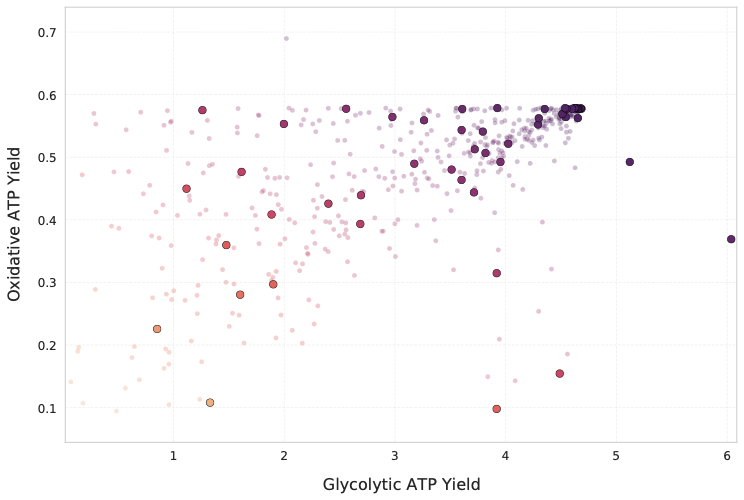

**Figure S6. Stratified sampling of kinetic models based on glycolytic and oxidative ATP yields of *BRCA1*^WT^ ovarian cancer.** Each dot represents one physiologically feasible steady state, colored by its sum of glycolytic and oxidative ATP production levels. Highlighted dots indicate the representative steady-state samples selected to capture distinct metabolic states for subsequent drug-response simulations.

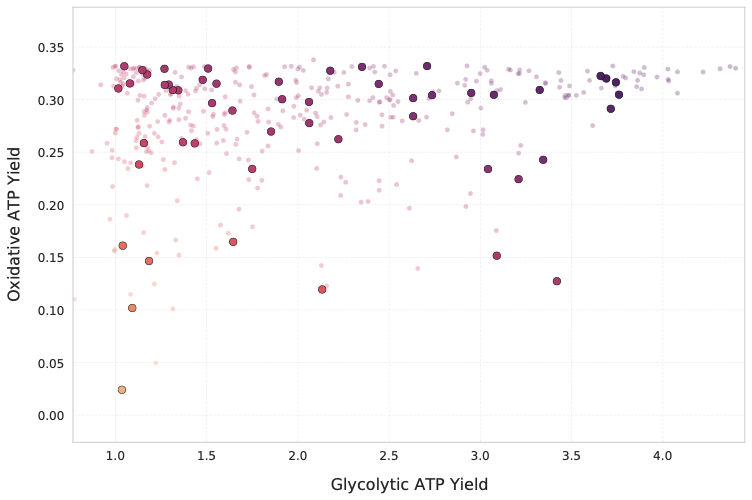

**Figure S7. Stratified sampling of kinetic models based on glycolytic and oxidative ATP yields of *BRCA1*^MUT^ ovarian cancer.** Each dot represents one physiologically feasible steady state, colored by its sum of glycolytic and oxidative ATP production levels. Highlighted dots indicate the representative steady-state samples selected to capture distinct metabolic states for subsequent drug-response simulations.

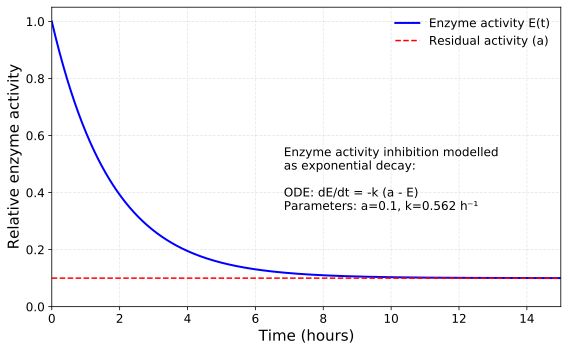

**Figure S8. Dynamics of enzymatic activity decay used to simulate drug inhibition.** The enzymatic activity decreases exponentially over time (blue line) toward the target inhibition level (red dashed line), representing the gradual onset of drug action during treatment simulations.

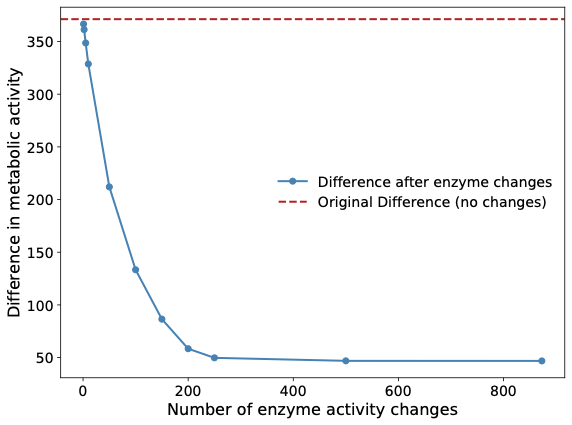

**Figure S9. Convergence of metabolic state alignment between *BRCA1*^MUT^ and *BRCA1*^WT^ physiologies as the number of allowed enzyme activity changes increases.** The residual flux deviation decreases sharply with the inclusion of up to 250 enzymes and plateaus, thereafter, indicating that a limited set of enzyme adjustments captures most of the metabolic differences between the two physiologies.

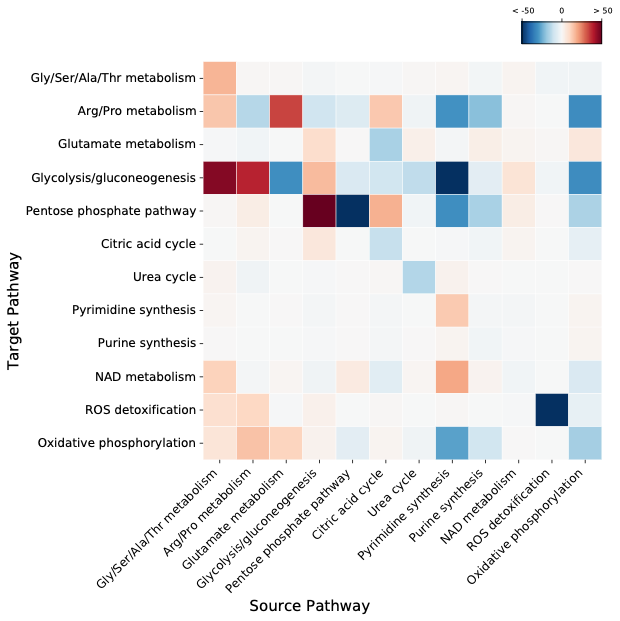

**Figure S10.** **Pathway effect scores capture the influence of enzyme perturbations from a source pathway to a target pathway.** The heatmap illustrates the magnitude of flux change in a target pathway, considering only enzyme perturbations from a single source pathway. The total effect on the fluxes of the target pathway can be positive (red) or negative (blue) based on the sign and magnitude of the enzyme fold changes from the source pathway. Terminal pathways like pyrimidine synthesis and Gly/Ser/Ala/Thr metabolism are regulated mainly by local changes within the pathway itself. In contrast, central carbon pathways like glycolysis and the pentose phosphate pathway are affected by multiple enzymes across different pathways, consistent with their role as the central hub of cellular metabolism.

**Table S1. Enzyme abbreviations and full names of predicted metabolic drug targets.**

| **Reaction ID** | **Name** |
| --- | --- |
| TRIOK | Triokinase |
| HMR_7748 | Alpha-D-Glucose 6-Phosphate Ketol-Isomerase |
| r0354 | Hexokinase |
| PGI | Glucose-6-Phosphate Isomerase |
| ICDHyrm | Isocitrate Dehydrogenase (NADP+) |
| ICDHxm | Isocitrate Dehydrogenase (NAD+) |
| r0426 | Isocitrate Hydro-Lyase |
| PPAP | Phosphatidic Acid Phosphatase |
| r0301 | Xanthosine-5-Phosphate:Ammonia Ligase (AMP-Forming) |
| 3DSPHR | 3-Dehydrosphinganine Reductase |
| SERPT | Serine C-Palmitoyltransferase |
| METAT | Methionine Adenosyltransferase |
| PSP_L | Phosphoserine Phosphatase (L-Serine) |
| r0178 | Succinate-Semialdehyde:NAD+ Oxidoreductase |
| r0179 | Succinate-Semialdehyde:NADP+ Oxidoreductase |
| TMDS | Thymidylate Synthase |
| NTD1 | 5'-Nucleotidase (dUMP) |
| DGK1 | Deoxyguanylate Kinase (dGMP:ATP) |
| ADSS | Adenylosuccinate Synthase |
| IMPD | IMP Dehydrogenase |
| r0474 | 2-Deoxyadenosine 5-Diphosphate:Oxidized-Thioredoxin 2-Oxidoreductase |
| URIK1 | Uridine Kinase (ATP:Uridine) |
| GMPS2 | GMP Synthase |
| HMR_4343 | Uracil Phosphoribosyltransferase |
| CYTK1 | Cytidylate Kinase (CMP) |
| UMPK2 | UMP Kinase (CTP) |
| MI1PP | Myo-Inositol 1-Phosphatase |
| Mcomplex1 | Mitochondrial complex I (NADH2_u10mi) |

**Table S2. Biomass precursor abbreviations and full names**

| **Metabolite ID** | **Name** |
| --- | --- |
| adp_c | Adenosine Diphosphate |
| ala_L_c | L-Alanine |
| arg_L_c | L-Arginine |
| asn_L_c | L-Asparagine |
| asp_L_c | L-Aspartate |
| atp_c | Adenosine Triphosphate |
| chsterol_c | Cholesterol |
| clpn_hs_c | Cardiolipin |
| ctp_c | Cytidine-5'-Triphosphate |
| cys_L_c | L-Cysteine |
| datp_n | Deoxyadenosine Triphosphate |
| dctp_n | Deoxycytidine-5'-Triphosphate |
| dgtp_n | Deoxyguanosine-5'-Triphosphate |
| dttp_n | Deoxythymidine-5'-Triphosphate |
| g6p_c | D-Glucose 6-Phosphate |
| gln_L_c | L-Glutamine |
| glu_L_c | L-Glutamate |
| gly_c | Glycine |
| gtp_c | Guanosine-5'-Triphosphate |
| h2o_c | Water |
| h_c | Proton |
| his_L_c | L-Histidine |
| ile_L_c | L-Isoleucine |
| leu_L_c | L-Leucine |
| lys_L_c | L-Lysine |
| met_L_c | L-Methionine |
| pail_hs_c | 1-Phosphatidyl-1D-Myo-Inositol |
| pchol_hs_c | Phosphatidylcholine |
| pe_hs_c | Phosphatidylethanolamine |
| pglyc_hs_c | Phosphatidylglycerol |
| phe_L_c | L-Phenylalanine |
| pi_c | Orthophosphate |
| pro_L_c | L-Proline |
| ps_hs_c | Phosphatidylserine |
| ser_L_c | L-Serine |
| sphmyln_hs_c | Sphingomyelin |
| thr_L_c | L-Threonine |
| trp_L_c | L-Tryptophan |
| tyr_L_c | L-Tyrosine |
| utp_c | Uridine-5'-TrIphosphate |
| val_L_c | L-Valine |
